# Sideways lipid presentation by the antigen-presenting molecule CD1c

**DOI:** 10.1101/2025.10.27.684910

**Authors:** Thinh-Phat Cao, Guan-Ru Liao, Tan-Yun Cheng, Yanqiong Chen, Laura Ciacchi, Thomas S. Fulford, Rachel Farquhar, Jade Kollmorgen, Jacob A. Mayfield, Adam P. Uldrich, Emily Zhi Qing Ng, Graham S. Ogg, Dale I. Godfrey, Nicholas A. Gherardin, Yi-Ling Chen, D. Branch Moody, Adam Shahine, Jamie Rossjohn

**Affiliations:** Infection and Immunity Program and Department of Biochemistry and Molecular Biology, Biomedicine Discovery Institute, Monash University, Clayton, Victoria 3800, Australia; Division of Rheumatology, Immunity and Inflammation, Brigham and Women’s Hospital, Harvard Medical School, Boston, MA 02115, USA; Department of Microbiology and Immunology, The Peter Doherty Institute for Infection and Immunity at the University of Melbourne, Melbourne, Victoria 3000, Australia; Chinese Academy of Medical Sciences Oxford Institute, Nuffield Department of Medicine, University of Oxford, Old Road Campus, Oxford OX3 7BN, UK; MRC Translational Immune Discovery Unit, Weatherall Institute for Molecular Medicine, University of Oxford, Headington, Oxford OX3 9DS, UK; Institute of Infection and Immunity, Cardiff University, School of Medicine, Heath Park, Cardiff CF14 4XN, UK

## Abstract

Here we report mass spectrometry analyses of endogenous lipids captured by CD1c when bound to an autoreactive αβTCR. CD1c bound twenty-six lipids with bulky headgroups that could not fit within the tight TCR-CD1c interface. We determined the crystal structures of CD1c presenting several gangliosides, revealing a general mechanism whereby two lipids, rather than one, are bound in the CD1c cleft. Bulky lipids were orientated sideways so that their polar headgroups protruded laterally through a side portal of the CD1c molecule - an evolutionarily conserved structural feature. The sideways presented ganglioside headgroups did not hinder TCR binding and so represent a mechanism that allows autoreactive TCR recognition of CD1c. In addition, *ex vivo* studies showed sideways presented gangliosides could also represent TCR recognition determinants. These findings reveal a general mechanism whereby CD1c simultaneously presents two lipid antigens from the top and side of its cleft that differs markedly from other antigen-presenting molecules.

---

αβ T cell activation relies on T cell receptor (TCR) contact with antigens bound to antigen-presenting molecules. While Major Histocompatibility Complex (MHC) molecules, the MHC class I related molecule (MR1), and CD1 molecules present peptides, metabolites, and lipids, respectively, the positioning of αβTCRs relative to antigen-presenting molecules is generally conserved(*1*). The membrane distal surface of the αβ heterodimer contacts the membrane distal surface of the antigen-presenting molecule, along with antigen protruding from the antigen-binding cleft(*2–5*). Accordingly, conventional depictions of this aligned, end-to-end contact mechanism show the antigen-presenting molecule below the αβTCR, and antigens protrude ‘upwards’ for direct αβTCR contact. For over three decades, this central end-to-end binding concept has shaped the general understanding of what governs αβTCR specificity for antigen and antigen-presenting molecules.

The other conserved feature of antigen display to T cells by MHC, MR1 and CD1 is 1:1 stoichiometry of antigen bound to antigen-presenting molecules(*6*). Recently, an exception was identified, where each human CD1b protein binds two lipid ligands, one on top of the other, so that the lower lipid acts as an inert scaffold and the upper lipid protrudes upward to act as the antigen contacting TCRs sitting ‘atop’ of CD1b(*7, 8*). Among the four human CD1 molecules, CD1c likely has unique immunological functions, as it is expressed on subsets of dendritic cells in lymphoid tissues, marks marginal zone B cells, and can be found on malignant lymphocytes(*9–11*). Yet among human CD1 antigen-presenting molecules, the least is known about the display mechanisms of CD1c(*12–14*). Taking advantage of a new lipidomics platform(*15*) to analyse cellular lipids trapped between CD1c and an αβTCR, we found that CD1c uses a distinct antigen-display mechanism, where it binds two orthogonally positioned lipids with distinct structural motifs that simultaneously bind and protrude through distinct portals on the top and side of the cleft to control TCR binding and response.

## RESULTS

### Trapping ligands between CD1c and the 3C8 TCR

Understanding of CD1c display to TCRs is limited (*16*). To understand self-lipid display, we sought to trap and analyse endogenous lipids in CD1c-lipid-3C8 TCR complexes. Secreted CD1c-lipid complexes from 293T cells and recombinant 3C8 TCR protein underwent size exclusion chromatography to separate CD1c-lipid complexes that do or do not bind the 3C8 TCR. After eluting lipids in organic solvents time-of-flight mass spectrometry (TOF-MS) detected lipids that promote or block TCR binding (*17, 18*). Recent implementation of sensitive MS methods now allow lipidome-scale detection(*15*), yielding complex patterns of lipid ligands with 630 unique ion chromatograms detected across 11 fractions, with each ‘molecular event’ corresponding to a lipid with defined by *m/z*, retention time and intensity values (**Figure 1**).

**Figure 1.**
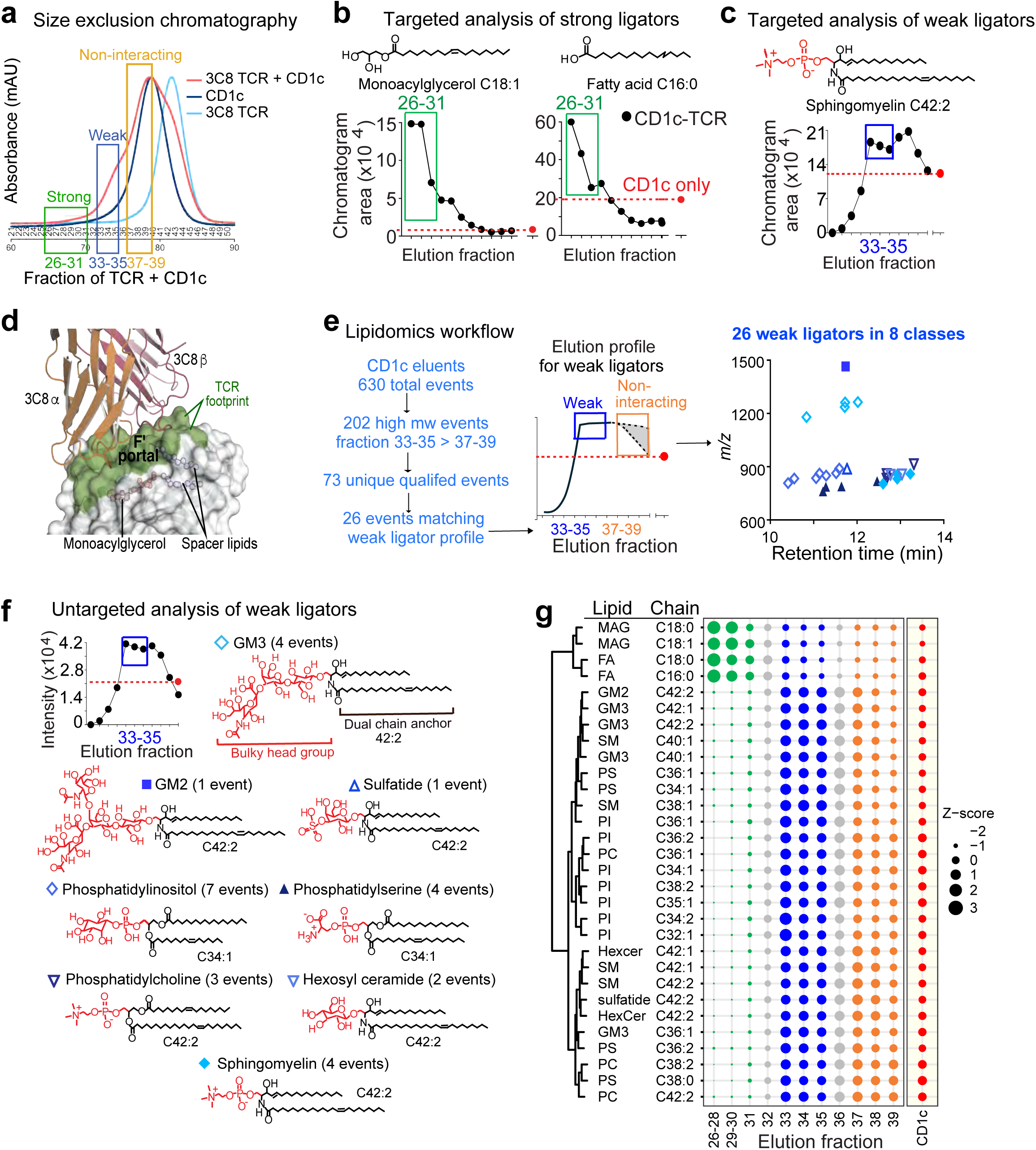
CD1c-TCR ligand trapping. **a.** Protein profiles of CD1c plus TCR show that complexes comigrating with CD1c monomers (fractions 37-39), and a biphasic curve with weakly (fractions 33- 35) and strongly (fractions 26-31) excluded complexes. **b-c.** Targeted MS analysis of lipids strongly and weaky excluded identified small monoacyl ligands, and a dual chain weak TCR ligator, sphingomyelin, respectively. **d, e, f.** Whereas the single chain headless lipid, monoacylglycerol (MAG), binds inside CD1c to allow a tight CD1c-TCR interface that surrounds the antigen exit portal (olive), lipidomics analysis of weak ligators of CD1c-TCR identified 26 dual chain lipids with large head groups (red), whose mechanism of CD1c binding was unknown. **g.** Unsupervised clustering analysis of ligands in strongly excluded fractions are headless single chain lipids, weakly excluded fractions show are dual chain lipids with large head groups. Elutional lipidomics was performed twice with similar results.

Fractions 37-39 of CD1c-TCR mixtures contained proteins that comigrated with separately added CD1c monomers, so these TCR-unbound CD1c proteins likely contained non-antigenic lipids (**Figure 1a**). Unexpectedly, early eluting CD1c-TCR complexes showed a biphasic response with strongly (fractions 26-31) and weakly (fractions 33-35) excluded complexes, which likely contained lipids that promoted durable versus intermittent CD1c binding to the TCR, respectively. To test this interpretation, we scanned fractions 26-31 for *m/z* values of two known antigens, C18:1 monoacylglycerol and C16:0 fatty acid(*16*), finding both, which directly demonstrated antigen trapping by CD1c-TCR (**Figure 1b**). Although weakly excluded fractions 33-35 were expected to contain weak T cell agonists, targeted analysis instead detected C42:2 sphingomyelin (42:2 SM), which is a known TCR blocker for CD1a(*19*) and CD1d(*20, 21*) (**Figure 1c**). This result was unexpected because prior work showed that the 3C8 TCR docked flush to CD1c, fully surrounding the F’-portal(*16*) (**Figure 1d**), which predicts that phosphocholine unit of 42:2 SM cannot fit between CD1c and the TCR, and it was known that shorter length SM (34:1) could act as a blocker of autoreactive CD1c-restricted TCRs (*16*).

### Lipidome-scale analysis

To address this contradiction, unbiased analysis sought additional lipids from weakly excluded CD1c fractions that might identify ligand motifs. From 630 total CD1c-associated events, we censored alternate adducts and isotopes and focused on events that peaked in weakly excluded fractions that showed mass greater than that of C34:1 SM (molecular weight 702) (**Figure 1e**), yielding 26 events (**Figure S1**) solved by collisional mass spectrometry (CID-MS) (**Figure S2**). After demonstrating a lack of non-specific binding to TCR alone and confirmation of peak capture in early excluded fractions (**Figures 1e and Figure S3**), this analysis detected dual chain lipids with bulky headgroups, including three additional SMs and twenty-two other lipids in seven classes: ganglioside GM3 (GM3), ganglioside GM2 (GM2), hexosylceramide (hexCer), sulfatide, phosphatidylserine (PS), phosphatidylcholine (PC), and phosphatidylinositol (PI) (**Figure 1f**).

### Bulky sphingolipids are permissive to TCR binding

Thus, lipidomics outcomes supported two unexpected general conclusions. First, strongly and weakly excluded complexes released lipids that clustered in two structural motifs (**Figure 1g**), single chain headless molecules versus dual chain bulky lipids, which implies two modes of lipid antigen capture or display by CD1c. Second, the tight interface between CD1c-3C8 TCR(*16*) (**Figure 1d**) was inconsistent with bulky headgroup positioning at the membrane distal surface of CD1c as predicted by general CD1 presentation models(*22, 23*). Further functional testing highlighted this conundrum, as treatment of CD1c tetramers with bulky gangliosides, GM3 or GD3, failed to suppress the autoreactive recognition of CD1c with 3C8 TCR transfected SCARB1-deficient HEK293T cells, unlike that of the antigen-specific CD1b-restricted TCR control (*23*) (**Figure 2a**). Also, surface plasmon resonance (SPR) found similar steady state affinities (K_D_) of 3C8 TCR towards CD1c with endogenous lipids (CD1c-endo), CD1c-MAG and CD1c-GD3, as though lipids with absent, small or large headgroups all allow CD1c to bind the 3C8 TCR similarly (**Figure 2b**).

**Figure 2.**
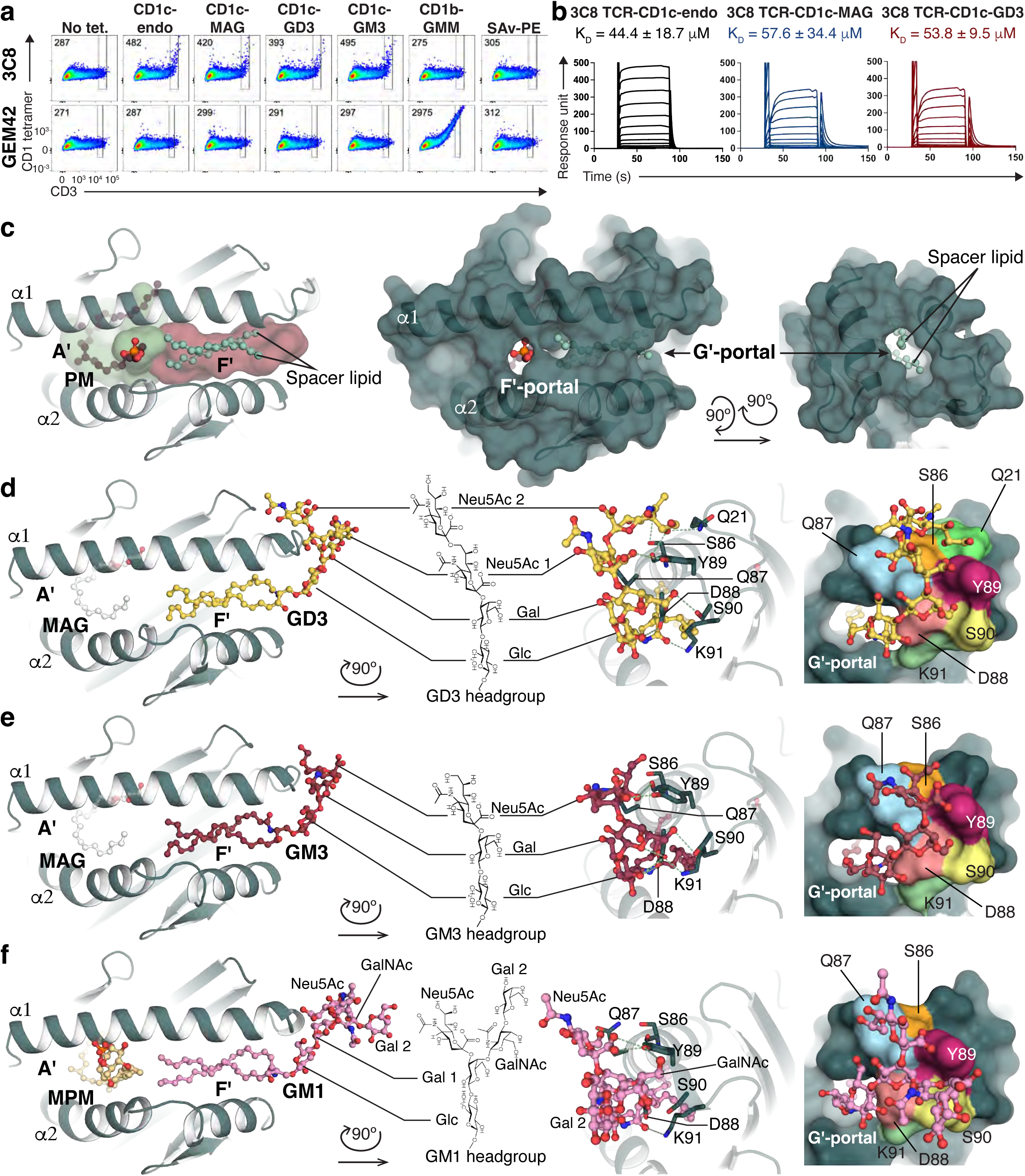
Two mechanisms of lipid antigen presentation by CD1c. **a.** FACS plots depicting HEK293T.SCARB1^-/-^ cells with surface expressed the 3C8 TCR (upper) and CD1b-restricted GEM42 TCR (lower) stained with CD1c tetramers with no added ligand (CD1c-endo), the indicated ligand or streptavidin-PE (SAv-PE) showing mean fluorescence intensities (MFI). **b.** Surface plasmon resonance (SPR) sensorgrams (n = 4 or 5) of the 3C8 TCR bound to the CD1c-lipid complexes show steady state affinities (K_D_). **c.** Left, Structure of CD1c presenting mycobacterial lipid PM. Right, surface representation demonstrates two major opening portals, F’ and G’. **d, e, f.** Structure of CD1c presenting gangliosides GD3, GM3, and GM1, respectively. Left, Top-down view showing MAG (white) or MPM (light yellow) in the A’-pocket, with GD3 (yellow), GM3 (red), or GM1 (pink) in the F’-pocket. Centre, Chemical structure of lipid headgroup and side view showing the headgroups protruding out of the G’-portal of CD1c. Details of interaction between sugars in the headgroups and CD1c are indicated, with hydrogen bonds represented as green dashes. Right, Surface representation showing the shape complementarity of the protein with the headgroups, with CD1c amino acids contacting the ganglioside headgroups individually colour coded. Sugar abbreviations: Glc, glucose; Gal, galactose; Neu5Ac, N-acetylneuraminic acid, or sialic acid; GalNAc, N-acetyl-galactosamine.

### CD1c uses a sideways antigen-displaying mechanism

CD1c cleft architecture is defined by the A’ and F’-pockets that bind lipids and connect to the external solvent via D’/E’,G- and F’-portals (*13*). The F’-portal allows antigen protrusion to the top of CD1c and other CD1 isoforms for TCR contact(*14*). We crystallised foreign phosphomycoketide (PM) with CD1c (**Figure 2c, Figure S4a, and Table S1**)(*24*), which resided in the A’-pocket in an upright position, similar to the seating of MAG in prior structures, but PM did extend through the F’-portal of CD1c, analogous to that of the CD1c-mannosylphosphomycoketide (MPM) structure (*14*). The F’-pocket also harboured one or two single chain spacer lipids, hinting that CD1c might normally bind more than one lipid.

To investigate how lipids with bulky headgroups are permissive for 3C8 TCR binding, we determined the structures of mock-treated CD1c, CD1c-GD3, CD1c-GM3, CD1c-MPM-GM1, and CD1c-MPM-GD3 (see methods and **Table S1** and **Figure S4b**). GD3 and GM3 differ by an additional sialic acid moiety in GD3 (**Figure 2d-e**), whereas GM1 ganglioside matches the core structure of GM3, but contains two additional branched sugars (**Figure 2f**). In the CD1c-GD3 and CD1c-GM3 structures, unbiased electron density corresponding to MAG was clearly visible in the A’-pocket, and was positioned similarly to that observed in previous 3C8 TCR-CD1c-endo ternary complex(*16*), as well as MAG bound in the CD1c-mock structure (**Figure S4b**). MPM was built into the density in the A’-pocket of the CD1c-MPM-GM1 and CD1c-MPM-GD3 except for the flexible mannosyl headgroup as observed in other CD1c-mycoketide structures^24^.

Unlike other CD1 structures, where antigen headgroups protrude to solvent through the F’-portal (*25*), no electron density was observed at the F’-portal in the CD1c-ganglioside structures. Instead, continuous unbiased electron density was observed that extended laterally across the F’-pocket and exited via a sideways G’-portal of CD1c (**Figures 2c-f, Figure S4c-f**). Here, the two tails of the ceramide moiety (**Figure 2d-f**) adopted the previously observed position occupied by spacer lipids^12^ (**Figures 2b, S4a-b**). Namely, the two chains of extended laterally across the F’-pocket, where the amide linkage hydrogen bonded to Asp88 and Lys91 (**Figure 2d-f and Figure S4),** which anchored the sphingosine chain on top of the fatty acyl chain in all structures (**Figures 2d-f, Figure S4, right column**). The fatty acid was resolved up to 18 carbons in length. Therefore, with the C18 sphingosine chain, we modelled the ceramide tail length of 36:0.

The bulky headgroups of GD3 and GM3 protruded through the G’-portal (**Figure 2d-e**), where the electron density for all sugars suggested a stable interaction with the outer surface of CD1c (**Figure S4c-d, and Figure S4f**). The branched sialic acid moiety of GM1 was not well resolved however, suggesting less stable contact with CD1c (**Figure S4e**). The bent headgroups of GD3 and GM3 wrapped around the extended loop (His84 – Phe94) of the α1-helix of CD1c and made a series of polar-mediated contacts with each sugar moiety (**Figure 2d-e**). Here, the surface contour of the α1-helix exhibited high shape complementarity with the GD3 headgroup, where Gln87 and Tyr89 generate a concave curvature to position the galactose (Gal) and two neuraminic acids (Neu5Ac). GM3, which lacks the terminal Neu5Ac, formed a similar curvature, but without contacting Gln21 and Ser86 (**Figure 2d-e**). The branched GM1 headgroup did not fully engage with CD1c and formed hydrogen bonds via N-acetyl-galactosamine (GalNAc) to Tyr89 (**Figure 2f**). The sideways positioning of the gangliosides provided immediate structural insight into their permissive nature for 3C8 TCR binding, as they adopt an unprecedented display mechanism whereby the bulky headgroups protrude sideways from the G’-portal of CD1c.

### CD1c volume versus lipid length

After implementing criteria for demarcating G’-portal boundaries, new (**Figure 2**) and prior CD1c structures ^3,12,13,24^ provided data for volume estimates of the CD1c cleft (*8*). The revised consensus estimate from all structures was 2050 **±** 180 Å^3^, suggesting that CD1c accommodate ∼50-54 CH2 units (C50-54). This value is ∼1.4-fold larger than estimates for the CD1a and CD1d cleft (∼C38-39) (*8*), which normally bind one ligand with two alkyl chains (∼C38). Thus, the CD1c ∼40 % larger cleft volume in CD1c is consistent with dual ligand binding in general, and the measured volume approximately matches the overall length of one single chain plus one dual chain lipid, as observed in individual structures reported here (**Figure 2c-f**).

### The G**’**-portal is a unique feature to CD1c

CD1c possesses a distinctive binding pocket that exhibits a mismatch with lipid size, attributable to its flexible F’-pocket. Compared to other CD1 isoforms, and despite the similarity in the overall structure (**Figure S5**)(*12, 13*), the F’-pocket of CD1c is readily open, so that dual tail lipids can replace the two spacer lipids, and the large headgroup can protrude via the G’-portal (**Figure 2c and 3a**). The G’-portal is formed by the arrangement of a constellation of residues (**Figure 3a**). The two α-helices of CD1c was remodelled upon accommodating these sideways ligands (**Figure 3b**). Despite this, the dimensions of the G’-portal did not undergo major change, with its major axis (∼16.0 – 17.2 Å) and minor axis (∼10 – 10.6 Å) remaining constant (**Figure 3c**). CD1c structurally aligned closely with the other three human CD1 isoforms (**Figure S5b**). However, whereas the smaller side chains of Ser90, Ser139, Ser143 and Leu147 make the solvent accessible G’-portal for CD1c, the corresponding positions in the other CD1 isoforms contain more bulky residues that block this G’-portal site (**Figure 3d**). As such, the G’-portal is essentially an invariant and unique feature of CD1c. Amino acid sequence alignments of *CD1C* genes across 15 mammalian species (**Figure S6 and Table S2**) showed ∼98% sequence identity among primates and 28-69% identity in non-primates. Except equine CD1c, conservation of the small residues that form the G’-portal were present in all species. Thus, this lateral portal is likely conserved throughout evolution of placental mammals.

**Figure 3.**
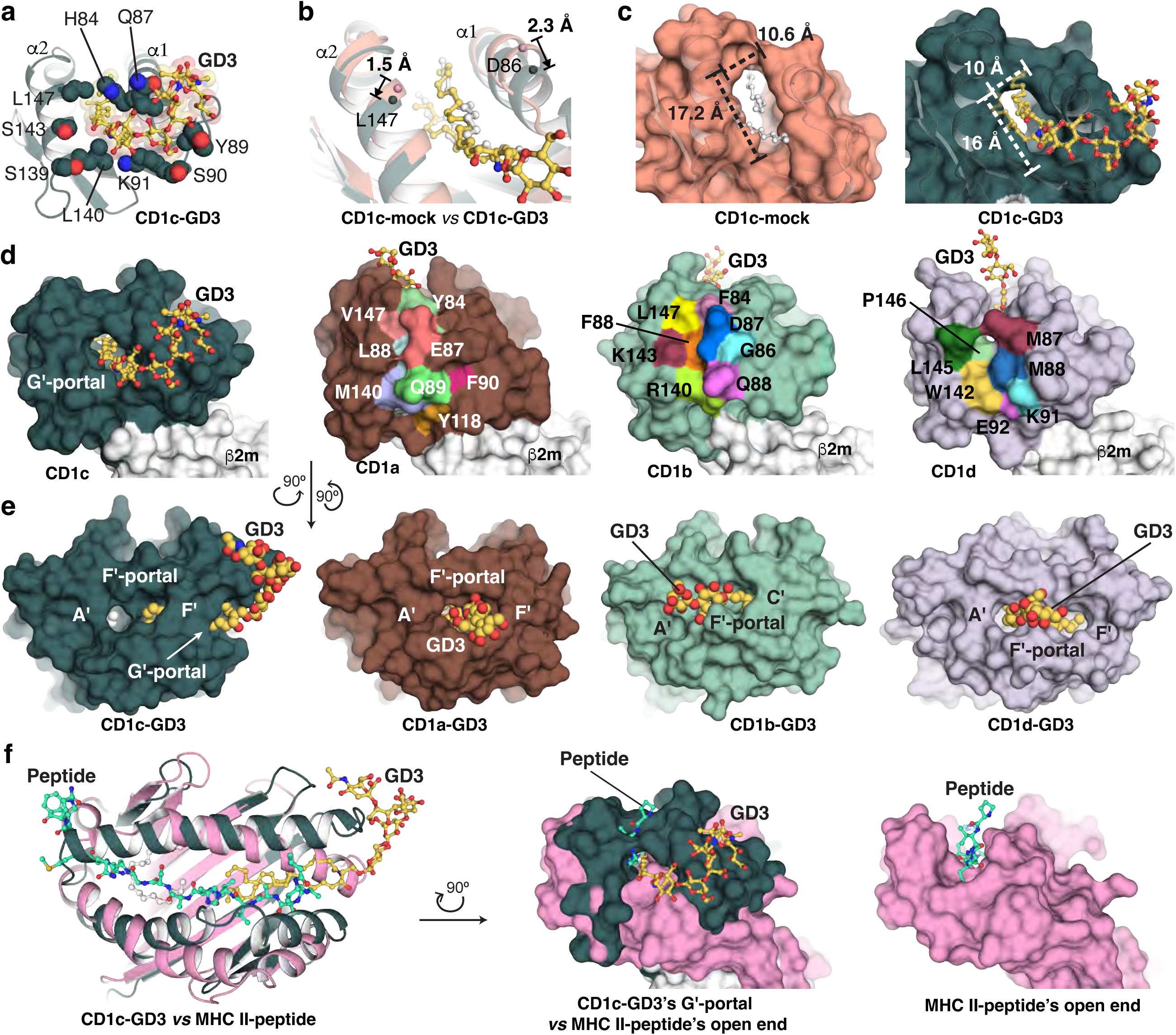
The G’-portal is a unique feature of CD1c. **a.** Details at the G’-portal of CD1c showing the protrusion of the GD3 headgroup (yellow ball-and-sticks). Amino acid side chains are depicted in space-filling format. The G’-portal is constituted by nine amino acids, including four large residues His84, Gln87, Tyr89, and Lys91, and five smaller residues that create the accessible entry point, i.e. Ser90, Leu140, Ser139, Ser143, and Leu147. **b.** Structural alignment of CD1c-GD3 (teal) versus CD1c-mock (pink) showing the movements of the a-helices, with arrows depicting direction of motion from the CD1c-mock to CD1c-GD3 structures. Ca atoms of Asp86 and Leu147 are depicted in spheres with the corresponding colours. Lipids are depicted in ball-and-stick, with GD3 coloured in yellow and spacer lipids in CD1c-mock coloured in white. **c.** Comparison of the G’-portal in the structures of CD1c-mock versus CD1c-GD3. The major axis is measured from Lys91 Cβ to Leu147 Cγ, and the minor axis is measured from His84 Cβ to Leu140 Cγ. **d.** Side view to the G’-portal of CD1c in comparison with the CD1a (brown), CD1b (green) and CD1d (blue), represented as surfaces. The blocking residues of the corresponding G’-portal position in each CD1 isoform are individually coloured as footprint. **e.** Top view to the F’-portal of all CD1 isoforms demonstrating the sideways presentation of CD1c while the same GD3 lipid (yellow spheres) is presented upright in the other isoforms. CD1d-GD3 was reported previously (PDB ID 3AU1) (*26*). **f.** Comparison of the CD1c G’-portal versus the open end of MHC II. CD1c and GD3 are in the same colour code as previous panels. The presented peptide in MHC II is depicted in green-cyan ball-and-sticks. Right, Equivalent position of the G’-portal in CD1c and the open end of MHC II. The structures are portrayed in surface representation. The structure of MHC II-peptide without superposition of CD1c-GD3 is placed at the right for reference. MHC II-peptide was reported previously (PDB ID 7T6I)(*33*).

To ascertain if sideways antigen presentation occurs with other CD1 isoforms, we determined the structures of CD1a-GD3 and CD1b-GD3 (**Table S1**) for comparison with the published CD1d-GD3 structure(*26*) (**Figures 3e and Figure S7**). For CD1a, CD1b and CD1d, the GD3 headgroup exited upwards through the F’-portal. Thus, the lateral protrusion mechanism is unique to CD1c, and can be compared to the open-ended cleft of MHC-II that allows overhang by the ragged ends of peptides(*6*). MHC-II has a notch that allows peptides to protrude upward, whereas by comparison the ganglioside headgroups escape sideways between the α1 and α2 helices of CD1c (**Figure 3f**). Both structural modifications likely exist as sizing mechanisms; the MHC-II allows for escape of excess peptide, but glycans egressing from the G’-portal adhere to the outside of CD1c and approach the TCR recognition site, so might tune T cell responses.

### Sideways presentation modulates recognition of upright lipids

The autoreactive 3C8 TCR interacted exclusively with CD1c, without co-contacting the headless MAG sequestered within the A’-pocket(*16*). To establish whether the sideway-presented GD3 ganglioside influences TCR recognition of a headed lipid in the A’-pocket of CD1c, we focused on the DN6 TCR, which demonstrates specificity for CD1c presenting the mycobacterial PM, and is predicted to dock over the F’-portal of CD1c(*27, 28*). The crystal structure of CD1c-PM (**Table S1**) confirmed the upward pointing nature of PM from the F’-portal (**Figure 2c**). SPR analysis demonstrated that the DN6 TCR remained similarly reactive to CD1c complexes treated with PM and GD3, compared to PM alone, but the response units (Rmax) was two-fold diminished compared to CD1c presenting PM alone (**Figure 4a**). Similarly, SPR analysis of another PM-specific TCR called 22.5(*28*) showed that, despite the same affinity for CD1c-PM and CD1c-PM-GD3, response units were ∼3-fold lower with CD1c-PM-GD3 GD3 (**Figure 4a**), suggesting that a sideways lipid impacted TCR binding in some manner. Similarly, after transducing the DN6 TCR into Jurkat cells, Jurkat.DN6 bound CD1c-PM tetramers in the presence and absence of GD3, but not to CD1c-endo, and CD1c-GD3 (**Figures 4b-c**), confirms the specificity of the T cell line for CD1c-PM and suggesting that the F’-sideway-presented GD3 still allowed the TCR interaction with the A’-upright-presented PM. However, consistent with the SPR data, the CD1c-PM-GD3 exhibited reduced staining on Jurkat.DN6 TCR cells.

**Figure 4.**
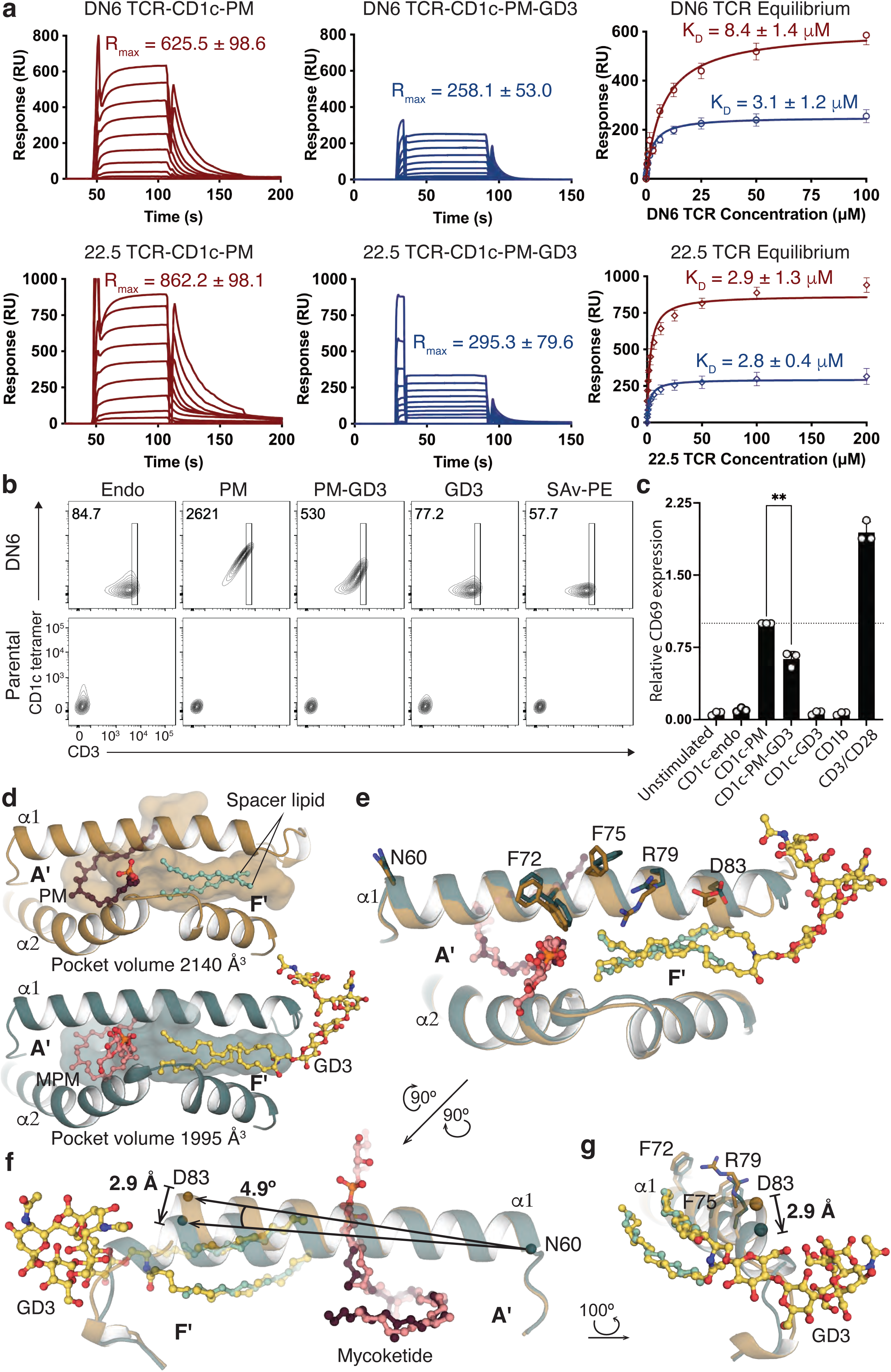
Sideways lipid presentation modulates TCR recognition of A’-upright lipids in CD1c. **a.** Steady state binding sensorgrams and equilibrium plots of DN6 (upper) and 22.5 (lower) TCRs against CD1c-PM with (blue) and without (red) the presence of GD3 using S{R. Error bars indicate mean ± S.D. of n=3 independent experiments, with respective steady state affinities (K_D_) and R_max_ values indicated in the equilibrium and sensorgram plots respectively. **b.** Staining of DN6 TCR transduced SKW-3 cells against CD1c-endo, -PM, -PM-GD3, and -GD3 tetramers, with SAv-PE as negative control. Gate in black box indicates tetramer-positive cells, with corresponding mean fluorescence intensities (MFI) indicated in each plot. **c.** CD69 activation assay of DN6 transduced SKW-3 cells against plate-bound CD1c-lipids. Data points for n=3 independent experiments are indicated. t-test statistical analysis between CD1c-PM and CD1c-PM/GD3, with a p value < 0.005 (**). **d.** Binding pocket comparison of CD1c-PM (copper-yellow, open conformation) and CD1c-MPM-GD3 (deep teal, closed conformation). **e.** Structural alignment of CD1c-PM (open conformation) and CD1c-MPM-GD3 (closed conformation) showing the steadiness of the α2-helix and the movement of the α1 helix. Colour code is same as in **d**. **f.** Back view to the α1-helix showing the pivoting of the helix at Asn60 with and without the presence of GD3 ganglioside. Colour code is same as in **d**. Cα of Asn60 and Asp83 are depicted in spheres. **g.** Details of the movement of Asp83 (spheres) in the presence of GD3 ganglioside which leading to the displacement of Phe72, Phe75, and Arg79 on the α1-helix.

To investigate the mechanism, superposition of CD1c-PM and CD1c-MPM-GD3 demonstrated interchangeable ligand-dependent conformational states that represent the “open” and “closed” conformers. In the “open” CD1c-PM conformation the CD1c cleft volume was 2140 Å^3^, while the closed cleft volume of CD1c-MPM-GD3 was reduced to 1995 Å^3^ (**Figure 4d**). While the α2-helix remained relatively fixed (**Figure 4e**), the α1-helix in the closed form pivoted 4.9°, as the GD3 headgroup brought the C-terminus of the α1-helix downward by 2.9 Å (**Figure 4f**). This effect, in turn, displaced C-terminal region of the α1-helix that is critical for the binding of DN6 and 22.5 TCRs^28^ (**Figure 4g**). Thus, CD1c is a dynamic molecule, where the DN6 and 22.5 TCRs favour binding the open conformer, while GD3 promotes the closed conformer.

### CD1c-ganglioside mediated T cell responses

We next sought to investigate whether the sideways presented gangliosides represented TCR recognition determinants. Because CD1c tetramer^+^ cells are rare in healthy blood (*29*), and CD1c-endo tetramer^+^ cells were present at similar frequencies to CD1c-GD3 or CD1c-GM3 tetramer^+^ cells (**Figure 5a**), we FACS sort-enriched these cells from nine donors using GD3-loaded CD1c tetramers and expanded them *in vitro* prior to restaining with CD1c tetramers loaded with distinct lipids (**Figure 5b**). These expanded polyclonal T cells were enriched with CD1c tetramer^+^ cells, including those from both the αβ and γδ T cell lineages. Here, most T cells stained with both unloaded CD1c-endo tetramers as well as ganglioside-loaded CD1c tetramers suggesting that most CD1c-ganglioside-reactive clones exhibit direct CD1c autoreactivity, in line with the sideways-presented headgroup locating distally to the TCR docking platform. Nonetheless, staining profiles demonstrated that the staining intensity of some populations were indeed modulated by CD1c-ganglioside tetramers relative to CD1c-endo (**Figure 5b**).

**Figure 5.**
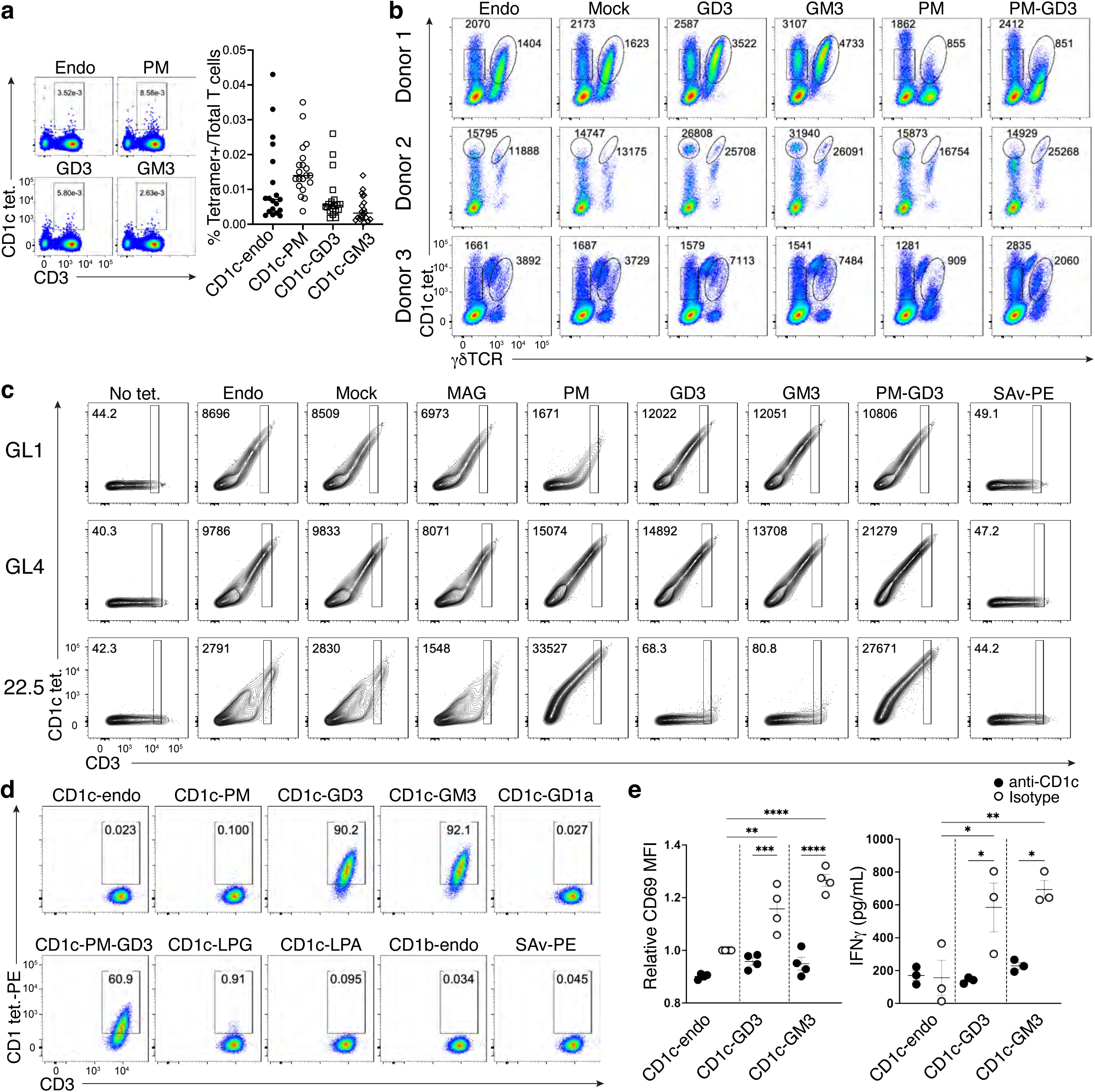
Autoreactive T cells towards CD1c presenting sideways gangliosides. **a.** (Left) Representative FACS plots showing CD1c tetramer staining of CD3^+^ T cells and CD3^-^ lymphocytes. Cells were stained with CD1c tetramers loaded with PM, GD3, or GM3, or with unloaded CD1c tetramers (CD1c-endo). (Right) Scatter plot showing frequencies of CD1c tetramer^+^ T cells within the T cell populations from *n* = 20 PBMC samples. **b.** Representative FACS plots showing *in vitro* expanded CD3^+^ T cells stained with CD1c tetramers loaded with the indicated lipids. The MFI of CD1c tetramers is shown next to each gate. **c.** Representative contour plots showing CD1-lipid tetramer staining on HEK293T.SCARB1^-/-^ cells transiently transfected to express CD1c restricted TCRs shown in **Figure S8b-c** or control TCRs, 22.5 (CD1c-PM reactive). MFI values of tetramer positive cells are shown in the top left of each plot, gated comparatively across all samples. The experiments were performed in two independent repeats. **d.** Representative FACS plots showing F10 T cell clone stained with CD1c tetramers loaded with the indicated lipids. **e.** CD69 expression and IFNγ production of F10 T cell clone after stimulation with bead-bound CD1c-lipid, which was pre-incubated with an anti-CD1c monoclonal antibody (0.5 μg/mL) or isotype control (0.5 μg/mL). Each symbol represents an independent repeat. CD69 expression data were normalised to relevant controls. Statistical analysis was performed using two-way ANOVA; *p < 0.05, **p < 0.01, ***p < 0.001, ****P<0.0001.

TCR sequencing of these candidate T cell clones revealed a polyclonal TCR repertoire, enriched for *TRBV4-1* for the αβ T cells and *TRDV1* for the γδ T cells, in line with previous reports (**Figure S8a**) (*29, 30*). TCRs from two such clones, one αβ (clone GL1) and one γδ (clone GL4) TCR, were sequenced and subsequently transfected for surface expression in SCARB1-deficient HEK293T cells and the cells subsequently stained with a panel of CD1c tetramers loaded with distinct lipids (**Figure 5c and Figure S8b-d**). CD1c-endo reactivity of the GL1 TCR was largely unaffected by the presence of gangliosides GD3 or GM3, however, staining was reduced by PM. This staining was rescued by CD1ctetramers loaded with both PM and GD3, suggesting that the presence of the ganglioside may fine tune PM-responsiveness. The GL4 TCR was largely unaffected when CD1c was loaded with distinct individual lipids, suggesting that the sideways-presented gangliosides are permissive to CD1c-autoreactivity. When tetramers were loaded with both PM and GD3, however, the staining intensity was approximately doubled relative to CD1c-endo, thereby suggesting that the co-presentation of PM an GD3 may represent a unique molecular determinant for antigenicity.

We next examined CD1c-ganglioside tetramer reactivity in tonsils, which are enriched in CD1c^+^ APCs compared to blood(*31, 32*). From three tonsil samples, dual CD1c-endo and CD1c-GD3 tetramer staining identified four αβ T cell lines with preferred reactivity towards CD1c-GD3 over CD1c-endo (**Figure S8e**). We sorted and expanded one CD4^+^ T cell clone (F10) (**Figure S8f)**, which confirmed greater reactivity towards CD1c-GM3 and CD1c-GD3 over CD1c-endo, and then extended these analyses to a broader range of CD1c-restricted ligands (**Figure 5d**). This confirmed lack of reactivity towards CD1c-endo and other ‘upward pointing’ antigens, including lysophosphatidylglycerol (LPG), lysophosphatidic acid (LPA) and PM, yet specific reactivity towards GD3 and GM3 (**Figure 5d**). As this F10 clone showed no reactivity towards GD1a, a branched ganglioside with two additional sugar headgroup moieties compared to GD3, this indicated that specificity determinants common to GD3/GM3 underlies this TCR-mediated recognition. The CD1c-ligand tetramer reactivity of the F10 T cell clone was also mirrored in activation assays, as measured by CD69 upregulation and IFN-γ production, which were blocked by an anti-CD1c mAb, thereby underscoring the CD1c-restricted nature of this T cell response (**Figure 5e**). Accordingly, while the sideways presentation of gangliosides can lead to the formation of TCR recognition determinants, the major consequence of such presentation is to not impede the broader CD1c-restricted autoreactive TCR repertoire.

## Discussion

Over four decades, structural studies on MHC-peptide, CD1-lipid and MR1-metabolite complexes provided foundational molecular insight into how antigens are displayed for T cell surveillance^1^. The generalist view is that antigens are anchored in 1:1 stoichiometry on antigen-presenting molecules and protrude upward toward TCR contact residues^8^. As such, it would not have been predicted that an antigen-presenting molecule would deviate from this central paradigm - the dual upwards and sideways presentation by CD1c was unexpected. The 2:1 lipid:CD1c stoichiometry and sideways presentation is likely a general mechanism for CD1c, as ganglioside loaded tetramers stain polyclonal T cells; simultaneous binding of one dual chain and one single chain lipid match the larger cleft volume of CD1c; and sequences that form the unique G’-portal are conserved across mammalian *CD1c* genes. These architectural features are not present in other CD1 isoforms, whereupon gangliosides are presented in the standard “upright” mode and are generally considered to represent inert antigens or block CD1-restricted T cell responses^11^.

What are the implications of CD1c presenting gangliosides, or other lipid-based antigens in a sideways manner? Firstly, direct T cell reactivity towards CD1c is a common mode of action for this isoform (*16, 29*) and our results suggest that, in general, the CD1c-restricted autoreactive T cells will not be impeded by ganglioside presentation. Secondly, the sideways presented ganglioside may represent a determinant for an innate receptor, or an atypical TCR binding site, the latter of which is supported by some CD1c-restricted T cell reactivity towards these antigens. Furthermore, γδ TCRs are known to decorate MHC-I like molecules in a range of docking modalities^33,34^, suggesting that sideways presented antigens might furnish specific ligands for γδ T cells. The findings reported here lay the foundation to establish whether CD1c-restricted T cell immunity involving gangliosides plays a role in homeostasis or disease where gangliosides are enriched, such as the nervous system or cancerous cells.

## Methods

### Protein expression and purification

Genes encoding human CD1c, CD1a, or hCD1d with human β2-microglobulin (β2m) were cloned into a single construct separated by a 2A self-cleavage peptide in the pHLSEC vector. The construct followed the sequence order of β2m-fos-2A-CD1c-jun, β2m-fos-2A-CD1a-jun, and β2m-fos-2A-hCD1d-jun with the fos/jun leucine zipper facilitating the post-translational association of the two proteins. Each gene was preceded by the mouse Igκ signal sequence to enable secretion of the heterodimer complex into the culture medium. The CD1 proteins was engineered with a hexa-histidine tag for affinity purification and an Avi™ tag for biotin labelling. Two thrombin cleavage sites were introduced before the fos/jun sequences. The single-plasmid construct encoding CD1c, CD1a, or CD1d and β2m was transfected into Expi293F GntI(-) cells. For CD1b, a similar design was constructed for insect *Trichoplusia ni* High Five cell expression. The secreted CD1c-β2m, CD1a-β2m, CD1b-β2m and hCD1d-β2m heterodimer complex (referred to hereafter as CD1c, CD1a, CD1b, and hCD1d) was harvested and subjected to a series of purification steps, including affinity chromatography, size-exclusion chromatography, and anion exchange chromatography, with purification validated by SDS-PAGE.

Genes encoding the α-and β-chains of DN6, 22.5, and 3C8 TCRs were cloned into the pET30a vector. For biotin labelling, the Avi™ tag was added to the β-chain of DN6 TCR and the α-chain of 3C8 TCR. These constructs were transformed into BL21 (DE3) *E. coli* cells. Protein expression was induced using 1 mM isopropyl β-D-1-thiogalactopyranoside, and expressed for four hours at 37°C before harvesting by centrifugation. The inclusion bodies were collected and washed with a buffer containing 50 mM Tris, 100 mM NaCl, 1% (v/v) Triton X-100, 2 mM dithiothreitol (DTT), 1 mM EDTA, and 0.5 mM phenylmethylsulfonyl fluoride at pH 8.0. The washed inclusion bodies were solubilized in a buffer comprising 50 mM Tris pH 8.0, 6 M guanidine hydrochloride, and 10 mM DTT. For refolding, 50 mg of each cognate α- and β-chain were gradually injected to a refolding buffer containing 50 mM Tris, 5 M urea, 440 mM L-arginine, 4 mM EDTA, 0.5 mM PMSF, 2 mM reduced glutathione, and 0.2 mM oxidized glutathione at pH 8.0. Refolding was carried out at 4°C for 48 hours. The refolded αβ heterodimers of each TCR were purified through sequential DEAE-cellulose anion exchange, strong anion exchange (HiTrap Q HP), size-exclusion chromatography, and hydrophobic interaction chromatography, with purification validated by SDS-PAGE. To improve the production yield of the 3C8 TCR, gene encoded for its α and β chains were separately reconstructed into pHLSEC vector for mammalian expression wherein the constant domains of both chains were further engineered with seven mutations described by Thomas *et* al ({Thomas, 2019 #42}), with an extra mutation of Lys44Gln on the α chain to assist the stabilized pairing between two chains. The purification of mammalian expressed 3C8 TCR followed the same method with that of CD1c protein.

### TCR trapping of CD1c-lipid complexes

Recombinant 3C8 TCR (1.75 mg) was mixed with recombinant CD1c protein (1.75 mg), 3C8 TCR alone (100 µg) or CD1c protein alone (100 µg) in Tris buffered saline (TBS) pH 8.0 and run over Sephadex 200 16/600 size exclusion column at 0.5 mL/min, using an AKTApurifier FPLC (GE). Approximately 2 µg of protein from protein containing fractions, based on UV elution profile, were analysed by SDS-PAGE for migration of TCR and CD1c species. In a separate experiment, 3C8 TCR alone (400 µg), CD1c protein alone (300 µg) or 3C8 TCR (6.5 mg) mixed with CD1c (4.5 mg) were run over two Sephadex 200 16/600 columns connected in series, at 0.5 mL/min. Fractions were collected and analysed by SDS-PAGE as per previous experiment. TCR trap experiments were completed twice with similar results.

### Lipid elution and analysis

CD1c-lipid complexes were eluted of lipids with the Bligh and Dyer extraction (*34*) that was adapted to small scale protein preparations (*15*). After normalization to 10 µM protein, fractions 26-28 and 29-30 were pooled due to low quantities within individual fractions. The injection volume was 10 µL for a reversed-phase HPLC column using an Agilent Poroshell EC-C18 column (1.9-micron, 3 x 50 mm) with a guard column (3 x 5 mm, 2.7 µm) using an Agilent 1260 HPLC system connected with an Agilent 6530 Accurate-Mass ESI-QToF mass spectrometer to acquire positive and negative ion mode data at a flow rate was 0.15 ml/min and the a published gradient (*35*). Intermediate eluting fractions (33–35) containing lipids allowing weak CD1c and 3C8 TCR interaction were compared to the late fractions (37–39) enriched for non-interacting CD1c and TCR monomers using Mass Hunter (Agilent) and the R package (version 3.4.2) XCMS (*36*) for lipidomic peak analyses and *in house* designed software methods(*37*). Lipid intensities are expressed as area-under-the-curve for identified lipids and were scaled and grouped by hierarchical clustering using Ward’s minimum variance method. Analysis and visualization was performed in R.

### Lipid identification

CID-MS analysis was performed on an Agilent 6546 Accurate Mass Q-TOF instrument and carried-out with a collision energy of 15-80 V with the isolation width set to 1.3 *m*/*z* and comparing signals to recently generated lipidomic maps or to synthetic standards when necessary.

### *In vitro* lipid loading

Monoacyl glycerol 17:0 (MAG) was obtained from Sigma-Aldrich (SMB00506), while gangliosides GD3, GM3, and GM1 were sourced from Avanti (860060, 860058, and 860065, respectively). phosphomycoketide (PM) and mannosyl β1-phosphomycoketide (MPM) were generously provided by Adriaan Minnaard (Groningen University). Lipids were solubilized via sonication in a buffer containing 10 mM HEPES, 150 mM NaCl, pH 7.5, and 0.5% tyloxapol. For direct loading, CD1c protein containing endogenous lipids (CD1c-endo) was incubated with 0.5% (w/v) tyloxapol (Sigma) in a buffer containing 10 mM HEPES and 150 mM NaCl, pH 7.5, at room temperature for 16 hours. The tyloxapol-treated CD1c protein (CD1c-mock) was subsequently incubated with (PM, MPM) at a molar ratio of 1:10. For crystallization, CD1c-endo was incubated with MAG at a molar ratio of 1:5 in the same buffer with 0.5% tyloxapol at room temperature for 16 hours. The CD1c-MAG complex was then further incubated with the desired lipids (PM, GD3, GM3, GM1) at a molar ratio of 1:10 under the same conditions as the MAG displacement step. For optimization of GM3 loading, GM3 was mixed to the GD3-loaded CD1c at a molar ratio of 10:1. For GM1 loading yields were insufficient for crystallization, so MPM was then used as displacement medium before treatment GM1. As a control, GD3-loaded CD1c was also mixed with MPM with lipid-loaded CD1c samples validated using analytical MonoQ anion exchange chromatography and Phastgel iso-electric focusing (Cytiva). For CD1a and CD1b, GD3 loading used methods for CD1a-sphingomyelin 42:2^19^ and CD1b-GMM^41^ complex formation, respectively. For tetramer staining that strictly required uniform biotinylation, biotin-labelled CD1c-Endo was aliquoted into 5 μg vials at 20 μM. Subsequently, the desired lipid stocks (2000 μM) were added (200 μM), and the mixture incubated at room temperature for 16 hours and validated using iso-electric focusing.

### Structural determination of CD1c-lipid complexes by X-ray crystallography

To facilitate crystallization, the CD1c construct was engineered by replacing its α3 domain with that of CD1b, a method previously reported (*14*) and utilized in our recent study(*24*). The resulting chimeric construct was cloned into the pHLSEC vector, maintaining a design similar to the wild-type CD1c construct to enable comparable expression, purification, and lipid loading protocols as described above. After loading the desired lipids, the leucine zipper was cleaved from the CD1c_chimeric_-β2m heterodimer complex by thrombin. Final size-exclusion chromatography was performed in a buffer containing 10 mM Tris and 150 mM NaCl at pH 8.0. The purified proteins were concentrated to 5 mg/ml and subjected to crystallization using the hanging drop vapor diffusion method. The mother liquor contained 100 mM N-cyclohexyl-2-aminoethanesulfonic acid (CHES) at pH 9.4, 1.05 M trisodium citrate, and 25 mM triglycine. Crystals of CD1c-lipid complexes were cryoprotected using 50% (w/v) sodium malonate and flash-frozen in liquid nitrogen. Crystals of CD1a-GD3 and CD1b-GD3 were crystallised as previously described(*19, 38*).

X-ray diffraction data were collected at the MX2 beamline of the Australian Synchrotron, part of ANSTO, and made use of the Australian Cancer Research Foundation (ACRF) detector. Data were processed using XDS(*39*) for indexing, integration, and symmetry assignment, followed by scaling with Aimless(*40*). Molecular replacement was performed using Phaser-MR(*41*) with the high-resolution structure of CD1c-MPM1 (PDB ID: 7MX4) as the search model for CD1c data sets, CD1a-SM 42:2 (PDB ID: 7KP0) for CD1a-GD3 dataset, and CD1b-PC (PDB ID: 6D64) for CD1b-GD3. Structures were refined using Coot(*42*) and phenix.refine(*43*). The three-dimensional structures of CD1 with different lipid ligands were visualized in PyMol, with the β-sheet of CD1c serving as a reference for structural alignment.

### Surface plasmon resonance

Steady-state equilibrium affinity measurements for CD1c-lipids and TCRs were conducted at 25°C using a BIAcore T200 instrument (Cytiva). The experiments utilized a running buffer containing 10 mM Tris and 150 mM NaCl at pH 8.0. Biotinylated CD1c-lipids were immobilized on streptavidin sensor chips to achieve approximately 2000 response units (RU) per flow cell, with hCD1d-endo serving as the reference control. TCRs, serving as analytes, were flowed over the sensor chip at 5 µl/min for 60 seconds, with the final response normalized by subtracting the response of the reference control. Serial dilutions were injected, with maximal concentrations of 100, 150, and 200 mM for DN6, 3C8, and 22.5 TCR, respectively. Affinity values and sensorgram plots were analysed and generated using BIAevaluation and GraphPad Prism software.

### Human sample collection

Peripheral blood mononuclear cells (PBMCs) were obtained from healthy human donors from the Australian Red Cross Blood Service after approval from the University of Melbourne Human Ethics Committee (1035100). Tonsils were collected from recurrent tonsillitis patients who consented to the use of their tissue for research at the John Radcliffe Hospital, Oxford, United Kingdom. Our study was reviewed and approved by the Oxford Radcliffe Biobank (ORB) Tissue Access Committee to obtain pseudonymised tissue samples and associated clinical data from patients recruited under ORB All procedures were performed according to the Declaration of Helsinki guidelines.

### Tetramer staining

Biotinylated CD1c monomers were co-expressed with biotin ligase in Expi293F GntI(-) cells and purified and the desired lipids were loaded using the direct loading method described above. For control (CD1c-mock), an equivalent volume of 0.5% tyloxapol was added and incubated overnight at RT. The next day, monomers were tetramerized with streptavidin-PE (BD), and cells were typically stained with 1 μg/ml tetramers. Peripheral blood mononuclear cells (PBMCs) or tonsil mononuclear cells (MNCs) were first incubated with anti-CD36 antibody (5-271; BioLegend; 10 μg/ml, 15 min, RT), followed by incubation with tetramers for 30 min at RT. Subsequent staining with an antibody cocktail was carried out at 4 °C prior to acquisition on an LSRFortessa (BD) and analysis using FlowJo software. For transient TCR expression, full TCR sequences were synthesized and subcloned into the pMIG-II plasmid. SRB1 KO HEK293T cells were co-transfected with the TCR and CD3 expression plasmids using FuGene HD. At three-day post-transfection, cells – as well as SKW3 stable cell lines – were stained with the indicated antibodies and CD1 tetramers loaded with specific antigens. For functional assessment, CD1c tetramer-positive T cells from PBMCs were sorted into U-bottom 96-well plates pre-coated with anti-CD3 (OKT3; 10 μg/ml) and anti-CD28 (Clone 28.2; 2 μg/ml) and cultured in the presence of rHuIL-2 (200 U/ml), rHuIL-7 (50 ng/ml), and rHuIL-15 (5 ng/ml). Cells are cultured in the complete medium consisting of RPMI-1640 supplemented with 10% fetal bovine serum (FBS) (JRH Biosciences), penicillin (100 U/ml), streptomycin (100 μg/ml), GlutaMAX (2 mM), sodium pyruvate (1 mM), nonessential amino acids (0.1 mM), Hepes buffer (15 mM) (pH 7.2 to 7.5) (Invitrogen, Life Technologies), and 2-mercaptoethanol (50 μM, Sigma-Aldrich). After two days, cells were transferred to new wells and co-incubated with 75K irradiated CD1c K562 cells, allowing T cell proliferation for 10–14 days in complete RPMI medium supplemented with cytokines. CD1c tetramer-positive T cells from tonsil MNCs were sorted into U-bottom 96-well plates and cocultured with irradiated PBMCs, allowing T cell proliferation for 14-21 days in complete RPMI medium supplemented with 10% heat-inactivated human serum and rHuIL-2 (200 U/ml; BioLegend).

### Bead-based activation assay

Biotinylated wild-type (WT) CD1c monomers loaded with the desired lipids were incubated with coreceptor antibodies (Miltenyibiotec) and magnetic beads (ThermoFisher) overnight at RT. The next day, coated beads were incubated with 50K expanded T cells in the complete RPMI medium supplemented with rHuIL-2 (25 U/ml; BioLegend) and anti-CD11a antibody (2.5 μg/mL, HI111; BioLegend) for at least 16 hours. For CD1c blockade experiments, the beads were incubated in complete RPMI medium in the presence of anti-CD1c antibody (L161; BioLegend) or isotype control (Mouse IgG1, κ; BioLegend) at 0.5 μg/mL at RT for 1 hour. T cells were stained with the indicated antibodies at 4 °C. The supernatant was harvested and the concentration of IFN-γ was measured by ELISA; BioLegend).

### ELISA

IFN-γ ELISA(BioLegend) used capture antibodies diluted in coating buffer, which were incubated at room temperature for 2 hours. The plates were then washed with PBS containing 0.1% Tween-20 and blocked with assay diluent at room temperature for 1 hour. Next, diluted supernatant samples and cytokine standards were added 2 hours. Following wash with PBS-Tween-20, detection antibodies were added and incubated for 1 hour. Avidin conjugated with HRP was incubated for 30 minutes. The reaction was developed using TMB substrate solution (ThermoFisher), and the reaction was stopped by adding the stop solution (ThermoFisher). Absorbance was measured at 450 nm using a ClarioStar plus reader (BMG Labtech).

### Plate-bound CD69 activation assay

Purified CD1c-lipids were coated on flat bottom 96-well polystyrene plates (Corning) at 12.5 μg/mL in phosphate-buffered saline pH 7.4 (PBS) at 4°C overnight. Wells were washed once using PBS and PM lipid was added to the final concentration of 2 μM followed by an incubation at 37°C for 4 hours. Approximate 100K SKW-3.DN6 cells were cultured in the coated plate for 16 hours before surface staining. Staining with an antibody cocktail was carried out at 4 °C before acquisition on an LSRFortessa (BD) and analysis using FlowJo software in triplicate.

## ACKNOWLEDGEMENTS

We thank the staff at the Australian Synchrotron MX beamline for assistance with data collection, and the Monash Macromolecular Crystallization Platform. We thank Shin Yi Tin for technical assistance and Adriaan Minnaard for the PM and MPM reagents. We acknowledge the contribution to this study made by Christine Jesus, Eve Warner, David Maldonado-Perez and the rest of the team at the Oxford Centre for Histopathology Research and the Oxford Radcliffe Biobank, which are funded by the University of Oxford, the Oxford CRUK Cancer centre, the NIHR Oxford Biomedical Research Centre (BRC) (Molecular Diagnostics Theme/Multimodal Pathology Subtheme and the NIHR CRN Thames Valley network. This work was supported by NIH AR048632, AI 049313, AI 162584 (DBM), the Wellcome Trust Discovery Award (302585/Z/23/Z, DBM, GSO, JR), the Australian Research Council (DP210103064), an NHMRC program grant (1113293), NHMRC Emerging Leadership Award 2027104 (AS) and 2027058 (NG), NHMRC Investigator Awards to 2008981 (JR) 2008913 (DIG), the UK Medical Research Council (GSO, YLC) and the Chinese Academy of Medical Sciences (CAMS) Innovation Fund for Medical Science (CIFMS), China (2024-I2M-2-001-1, GSO, YLC).

## AUTHOR CONTRIBUTIONS

T-PC, G-RL contributed equally, experimental design and primary data acquisition, performed analyses, solved structures and CD1c tetramer studies; T-YC and JAM undertook mass spectrometry and bioinformatic analyses, YC, LC, RF solved CD1 binary structures, JK, EZQN assisted with research; TSF and APU undertook TCR trap protein chemistry experiments, GSO, DIG, NAG and Y-LC supervised cellular immunology based experiments, provided reagents and funding, DBM, AS, JR are joint senior and corresponding authors, conceptualised and supervised project, provided funding and resources, co-wrote the paper alongside T-PC. All authors edited the manuscript.

## COMPETING INTERESTS STATEMENT

The authors have intellectual property though the Massachusetts General Brigham (DBM, GSO, JR) and the University of Oxford (GSO, YLC). GSO and YLC have relevant research collaborations with J&J Innovation. DIG and NAG hold PCT patents on therapeutic manipulation of CD1 family members.

## DATA AVAILABILITY

The crystallographic datasets generated and analysed within the current study were deposited to the Protein Data Bank (PDB) under codes 9OHX (CD1c presenting endogenous lipids), 9OHY (CD1c presenting phosphomycoketide in its open conformation), 9OHV (CD1c presenting dual lipids MPM and GD3 ganglioside), 9OHW (CD1c presenting GM1 ganglioside), 9OHU (CD1c presenting GM3 ganglioside), 9OHT (CD1c presenting GD3 ganglioside), 9OHZ (Crystal structure of CD1a presenting ganglioside GD3), and 9OI0 (Crystal structure of CD1b presenting ganglioside GD3).

## CODE AVAILABILITY STATEMENT

Neither custom code nor mathematical algorithms were used for this study.

