## Supplementary material for "Sideways lipid presentation by the antigen-presenting molecule CD1c": supp info

### Supplementary data

### Supplementary figures 1-8

### Supplementary tables 1-2

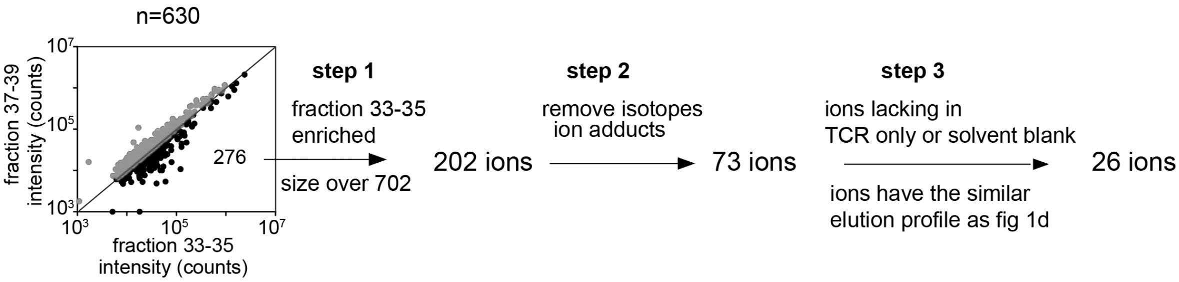


26 ions in 8 lipid classes

| | lipid class | detected *m/z* | formula | calculated *m/z* | lipid chain |
|---|---|---|---|---|---|
| 1 | phosphatidyl-serine (PS) | 760.5150 | C40H75NO10P [M-H]- | 760.5134 | 34:1 |
| 2 | | 786.5299 | C42H77NO10P [M-H]- | 786.5291 | 36:2 |
| 3 | | 788.5459 | C42H79NO10P [M-H]- | 788.5447 | 36:1 |
| 4 | | 818.5922 | C44H85NO10P [M-H]- | 818.5917 | 38:0 |
| 1 | sphingo-myelin (SM) | 803.6261 | C44H88N2O8P [M+HCOO]- | 803.6284 | 38:1 |
| 2 | | 831.6579 | C46H92N2O8P [M+HCOO]- | 831.6597 | 40:1 |
| 3 | | 857.6768 | C48H94N2O8P [M+HCOO]- | 857.6753 | 42:2 |
| 4 | | 859.6897 | C48H96N2O8P [M+HCOO]- | 859.6910 | 42:1 |
| 1 | phosphatidyl-inositol (PI) | 807.5039 | C41H76O13P [M-H]- | 807.5029 | 32:1 |
| 2 | | 833.5189 | C43H78O13P [M-H]- | 833.5186 | 34:2 |
| 3 | | 835.5350 | C43H80O13P [M-H]- | 835.5342 | 34:1 |
| 4 | | 849.5501 | C44H82O13P [M-H]- | 849.5499 | 35:1 |
| 5 | | 861.5501 | C45H82O13P [M-H]- | 861.5499 | 36:2 |
| 6 | | 863.5660 | C45H84O13P [M-H]- | 863.5655 | 36:1 |
| 7 | | 889.5810 | C47H86O13P [M-H]- | 889.5812 | 38:2 |
| 1 | phosphatidyl-choline (PC) | 832.6076 | C45H87NO10P [M+HCOO]- | 832.6073 | 36:1 |
| 2 | | 858.6199 | C47H89NO10P [M+HCOO]- | 858.6230 | 38:2 |
| 3 | | 914.6839 | C51H97NO10P [M+HCOO]- | 914.6856 | 42:2 |
| 1 | hexosylceramide (hexcer) | 854.6723 | C49H92NO10 [M+HCOO]- | 854.6727 | 42:2 |
| 2 | | 856.6880 | C49H94NO10 [M+HCOO]- | 856.6883 | 42:1 |
| 1 | sulfatide | 888.6228 | C48H90NO11S [M-H]- | 888.6240 | 42:2 |
| 1 | GM3 ganglioside | 1179.7337 | C59H107N2O21 [M-H]- | 1179.7372 | 36:1 |
| 2 | | 1235.7961 | C63H115N2O21 [M-H]- | 1235.7998 | 40:1 |
| 3 | | 1261.8131 | C65H117N2O21 [M-H]- | 1261.8154 | 42:2 |
| 4 | | 1263.8283 | C65H119N2O21 [M-H]- | 1263.8311 | 42:1 |
| 1 | GM2 ganglioside | 1464.8962 | C73H130N3O26 [M-H]- | 1464.8948 | 42:2 |

**Figure S1. Untargeted lipidomics analysis of weakly ligating lipids.** From 630 total mass spectral events, 276 show higher signals in the intermediate fractions 33-35 formed from complexes with weak CD1c-TCR exclusion, with 202 events with a mass larger than sphingomyelin (m/z 702). After removing redundant events derived from isotopes and alternate adducts, and solvent blanks, 26 event profiles that maximized in intermediate fractions were selected, from which 26 ions in 8 lipid classes were solved by matching to the mass of known self lipids and collisional mass spectrometry as detailed in Figure S2.

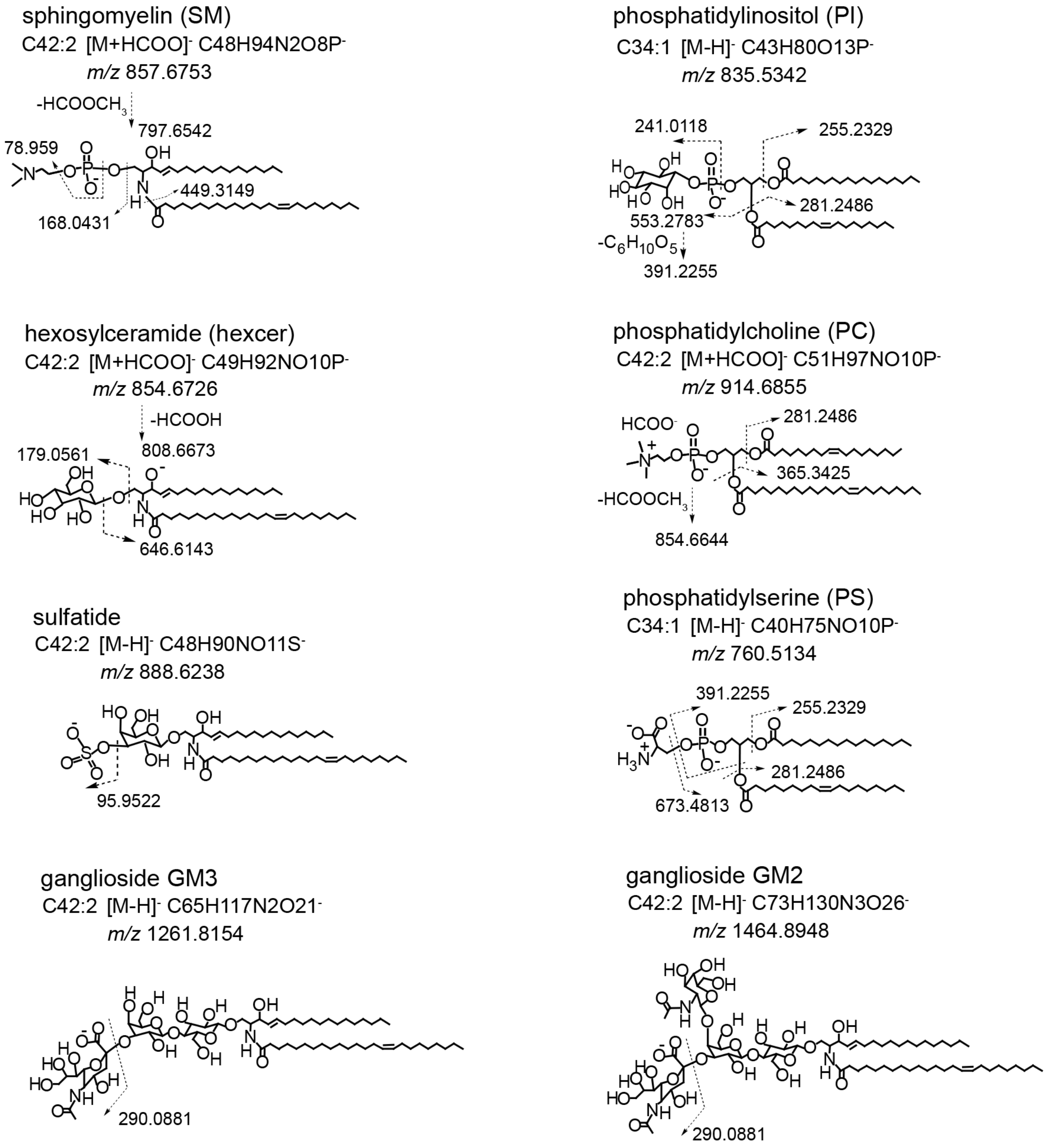


**Figure S2. Collisional mass spectrometry of lipids that generate weak size exclusion.**

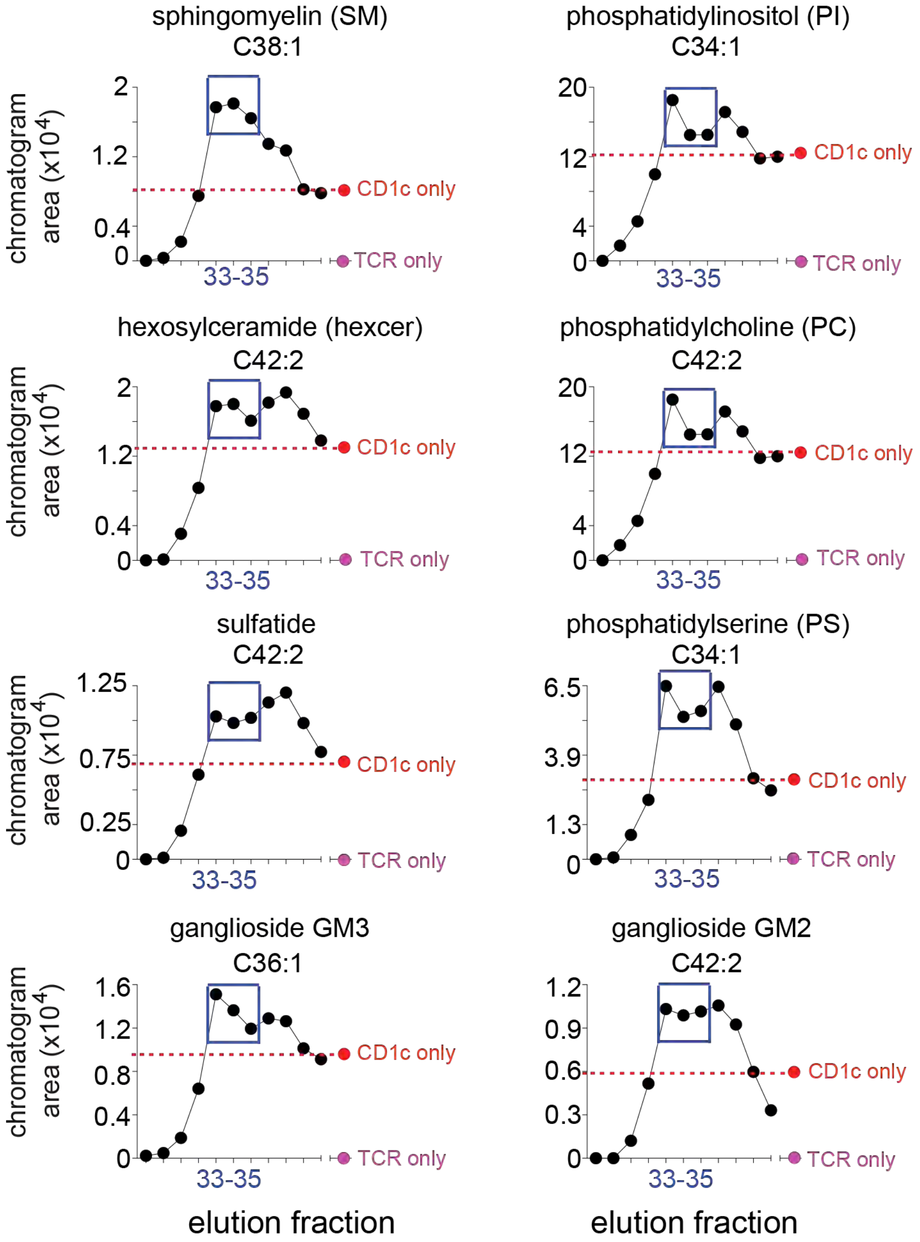


**Figure S3. Size exclusion chromatography elution profiles of complexes carrying lipids that generate weak exclusion patterns are shown for one lipid in each of 8 detected families.** These profile were selected for their peak signals in fractions 33-35 (blue box) that migrated just ahead of CD1c monomers. Signals from CD1c along (red) and TCR alone (pink) are superposed as positive and negative controls for lipid binding.

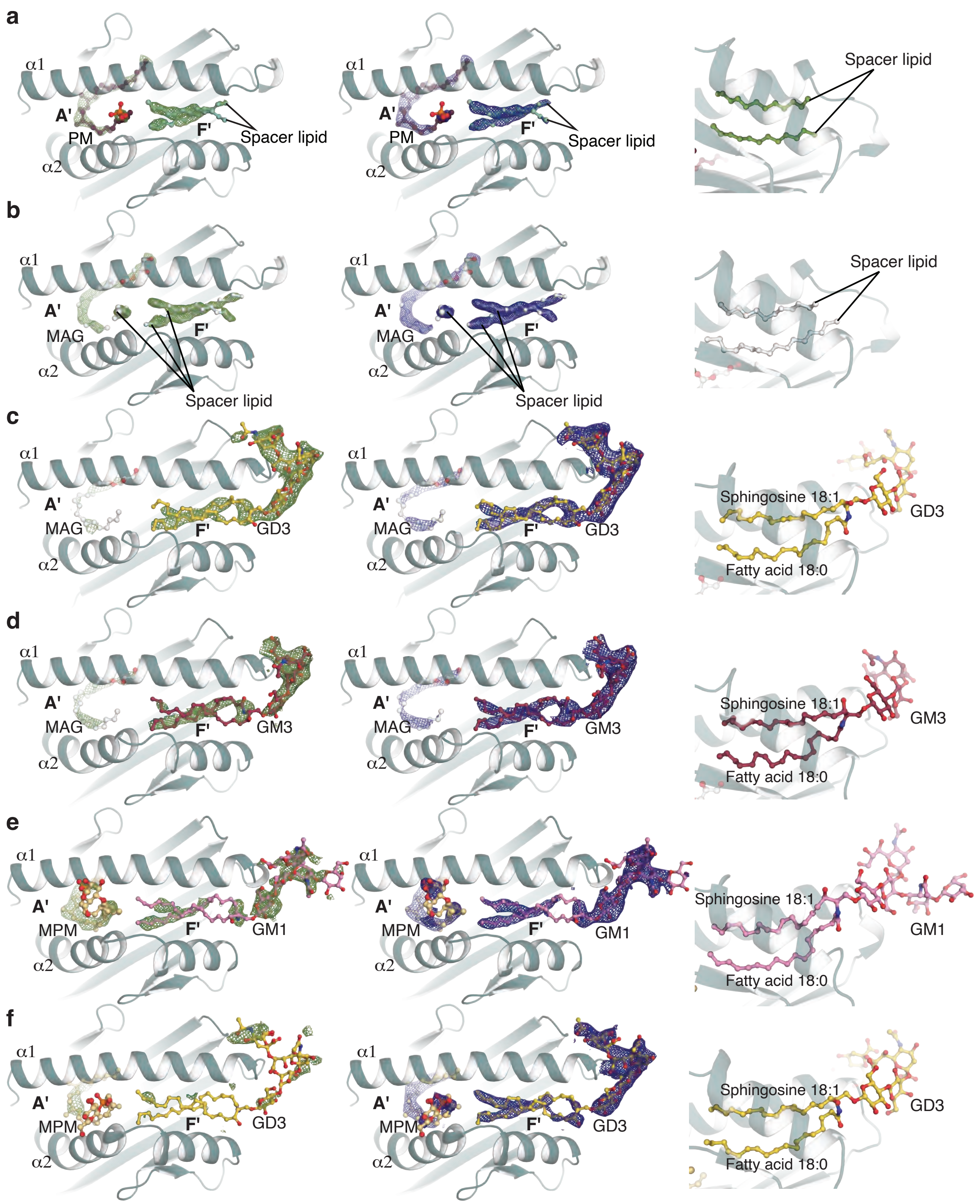


**Figure S4. Electron density of CD1c-PM, CD1c-mock, CD1c-gangliosides, and detail of lipids in F'-portal.**

Unbiased density (left, green) is contoured at 2.2σ, while refined density (middle, blue) is at 0.8σ. Structures are displayed as CD1c-PM, CD1c-mock, CD1c-GD3, CD1c-GM3, CD1c-GM1, and CD1c-MPM-GD3 in panels **a**, **b**, **c**, **d**, **e** and **f**, respectively. The right column shows the detail of ceramide tail in the F'-portal that replaces the position of the usual spacer lipids as seen in CD1c-PM and CD1c-mock.

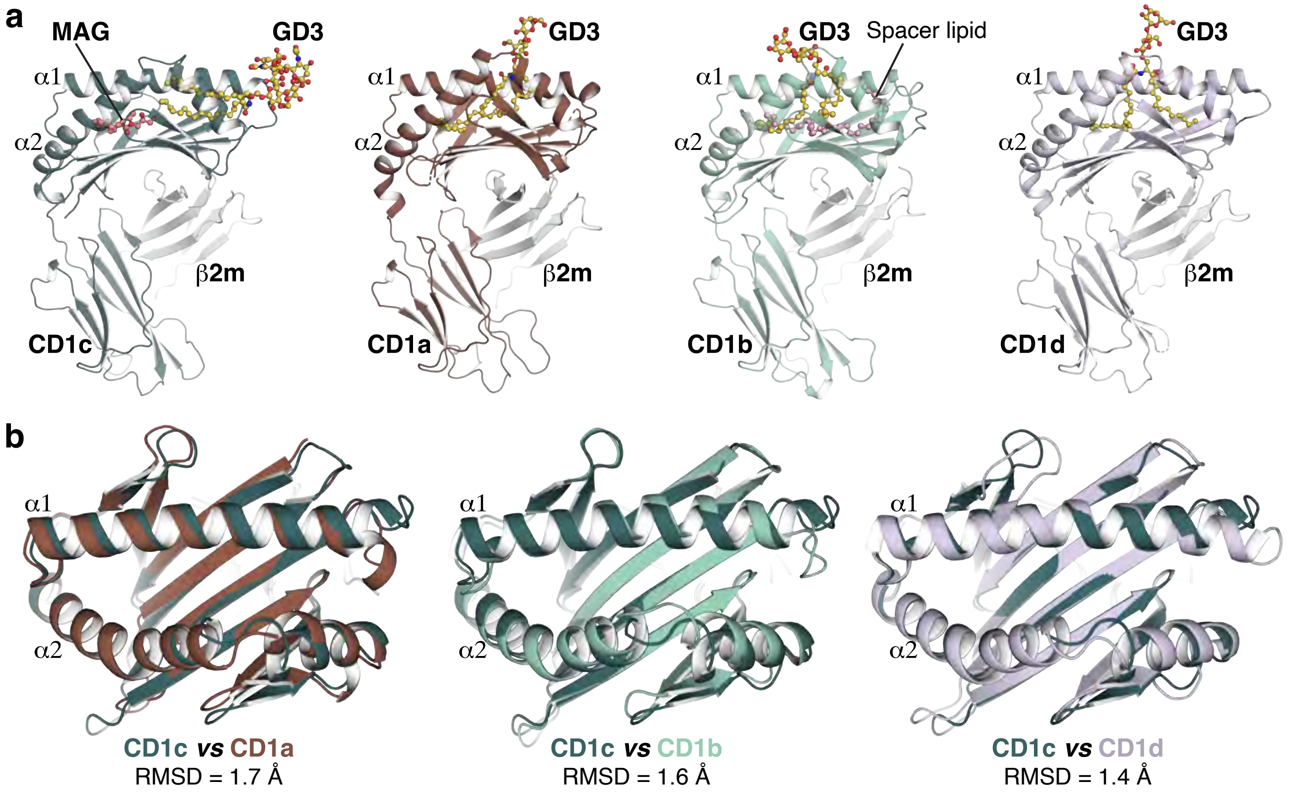


**Figure S5. Structural comparison of CD1c with CD1a, CD1b, and CD1d.**

**a.** Overall structure of CD1c-GD3 (teal, this study) in comparison with CD1a (brown, this study), CD1b (green, this study), and CD1d (blue, PDB ID: 3AU1). All the CD1 isoforms consist of a heavy chain associated with β2-microglobulin (β2m, white). **b.** Superposition in the binding pocket of CD1c versus CD1a, CD1b, and CD1d. The colour code is same as in panel **a**.

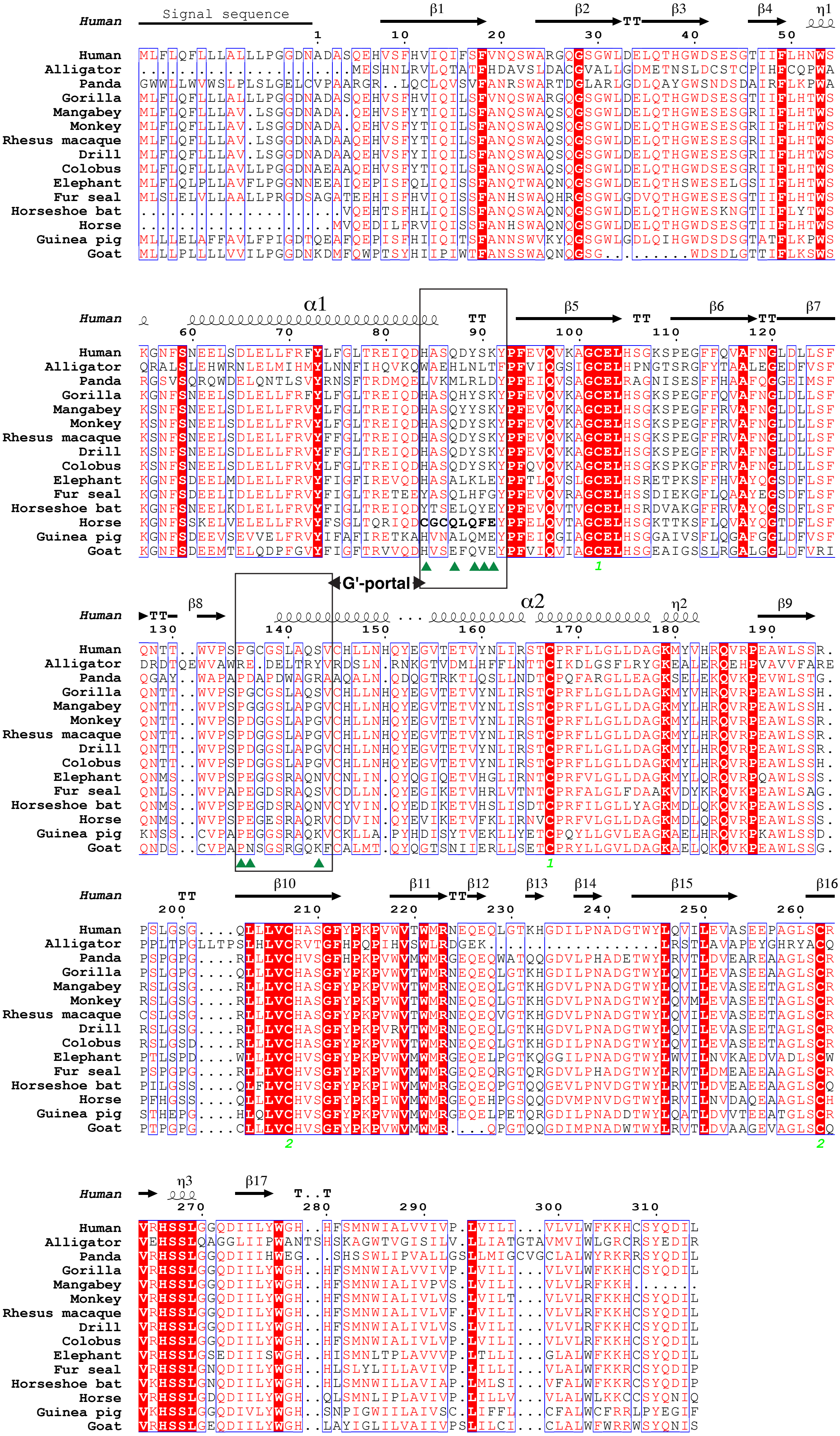

Signal sequence
β1
β2
β3
β4
η1
α1
β5
β6
β7
G'-portal
β8
α2
η2
β9
β10
β11
β12
β13
β14
β15
β16
η3
β17
Human
Alligator
Panda
Gorilla
Mangabey
Monkey
Rhesus macaque
Drill
Colobus
Elephant
Fur seal
Horseshoe bat
Horse
Guinea pig
Goat

**Figure S6. Primary sequence alignment of human CD1c versus CD1c from other fifteen representative species.**

Sequences of CD1c protein from other species were collected from Uniprot database and aligned using Muscle Alignment tool, then visualised with ESPript. Secondary structure assignment was based on the structure of CD1c-GD3, with $\alpha 1$ and $\alpha 2$ helices following the conventional assignment of CD1 molecule system. Residues that constitute the opening of the G'-portal are marked with green triangles within the black boxes, referred to human CD1c structure.

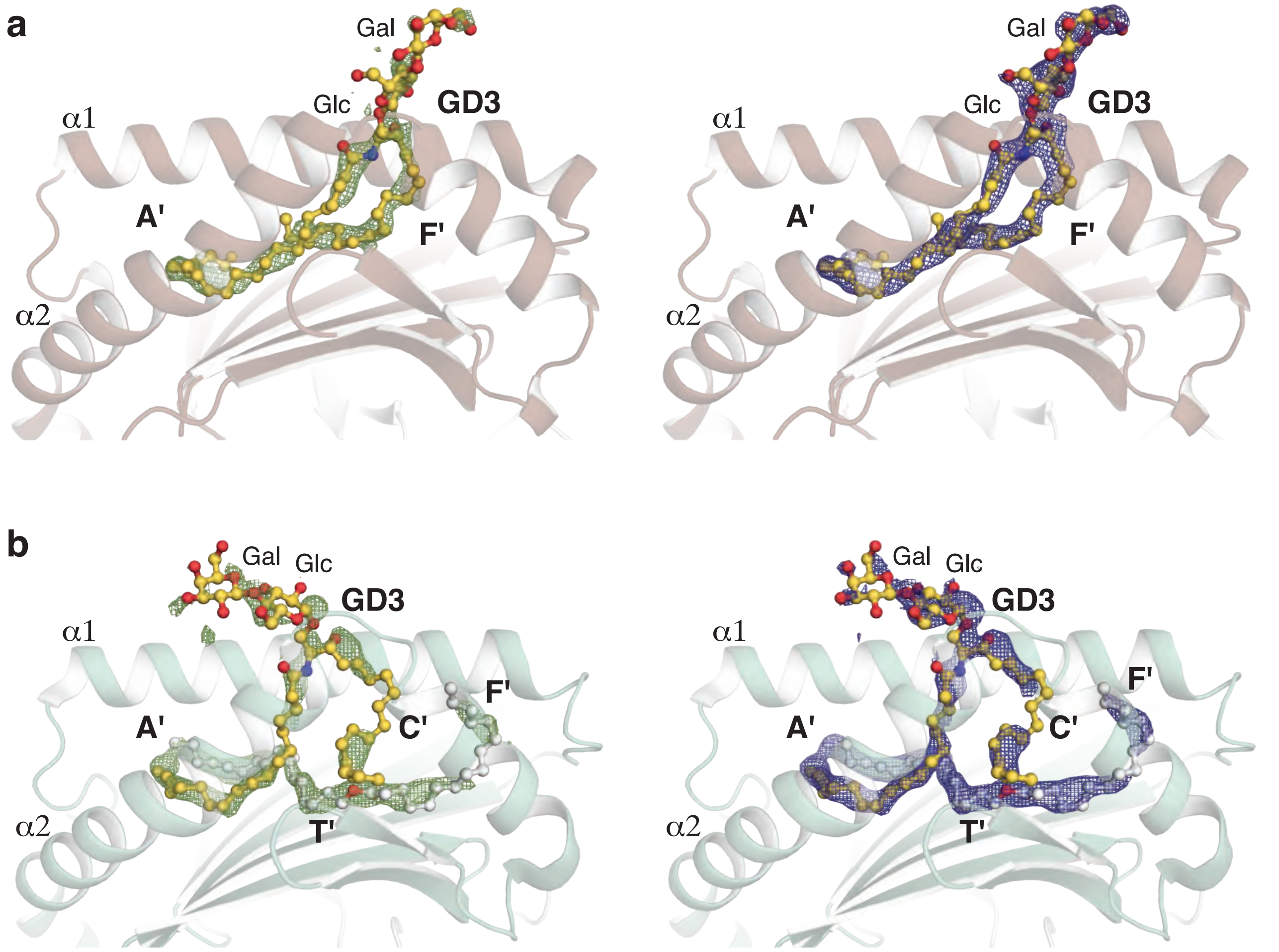


**Figure S7. Electron density of CD1a-GD3 and CD1b-GD3**

**a.** Structure of CD1a-GD3. The ceramide tail of GD3 is buried inside the A'- and F'-pocket. **b.** Structure of CD1b-GD3. The ceramide tail of GD3 is buried inside the A'- and C'-pocket. Unbiased density (left, green) is contoured at 2.2σ, while refined density (right blue) is at 0.8σ. The headgroup of GD3 in CD1a (red, **a**) and CD1b (green, **b**) is resolved up to two sugars, glucose (Glc) and galactose (Gal).

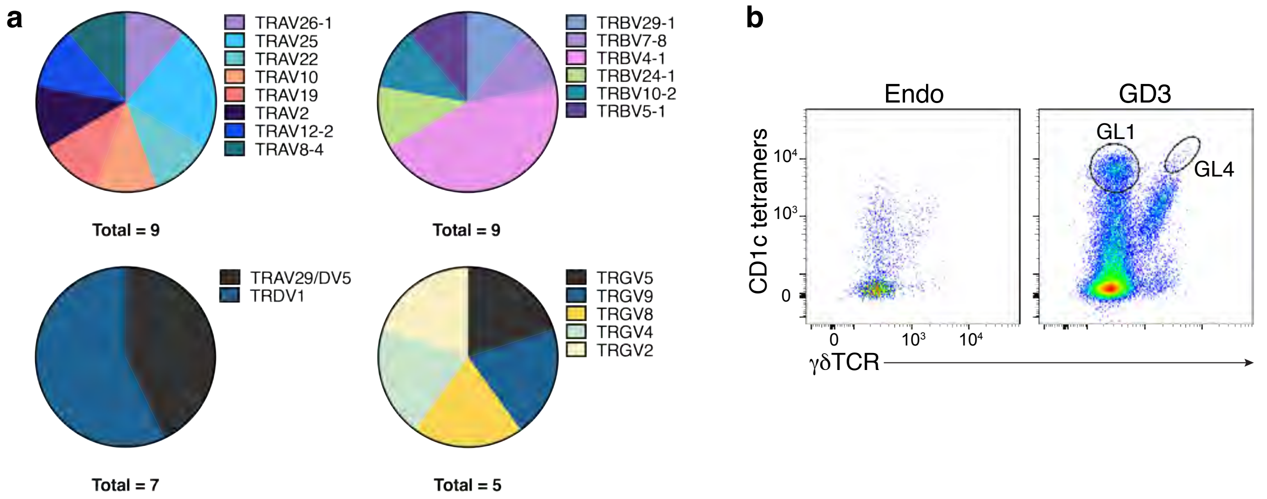


c

| Clone | TRAV/TRDV | CDR3α/δ | TRAJ/TRDJ | TRBV/TRGV | CDR3β/γ | TRBJ/TRGJ |
|---|---|---|---|---|---|---|
| GL1 | TRAV12-2 | CASYSGGGADGLTF | TRAJ45 | TRBV5-1 | CASSLSRNLYEQYF | TRBJ2-7 |
| GL4 | TRDV1 | CALGRVGPYKLIF | TRDJ1 | TRGV4 | CATWDGYKKLF | TRGJ1 |

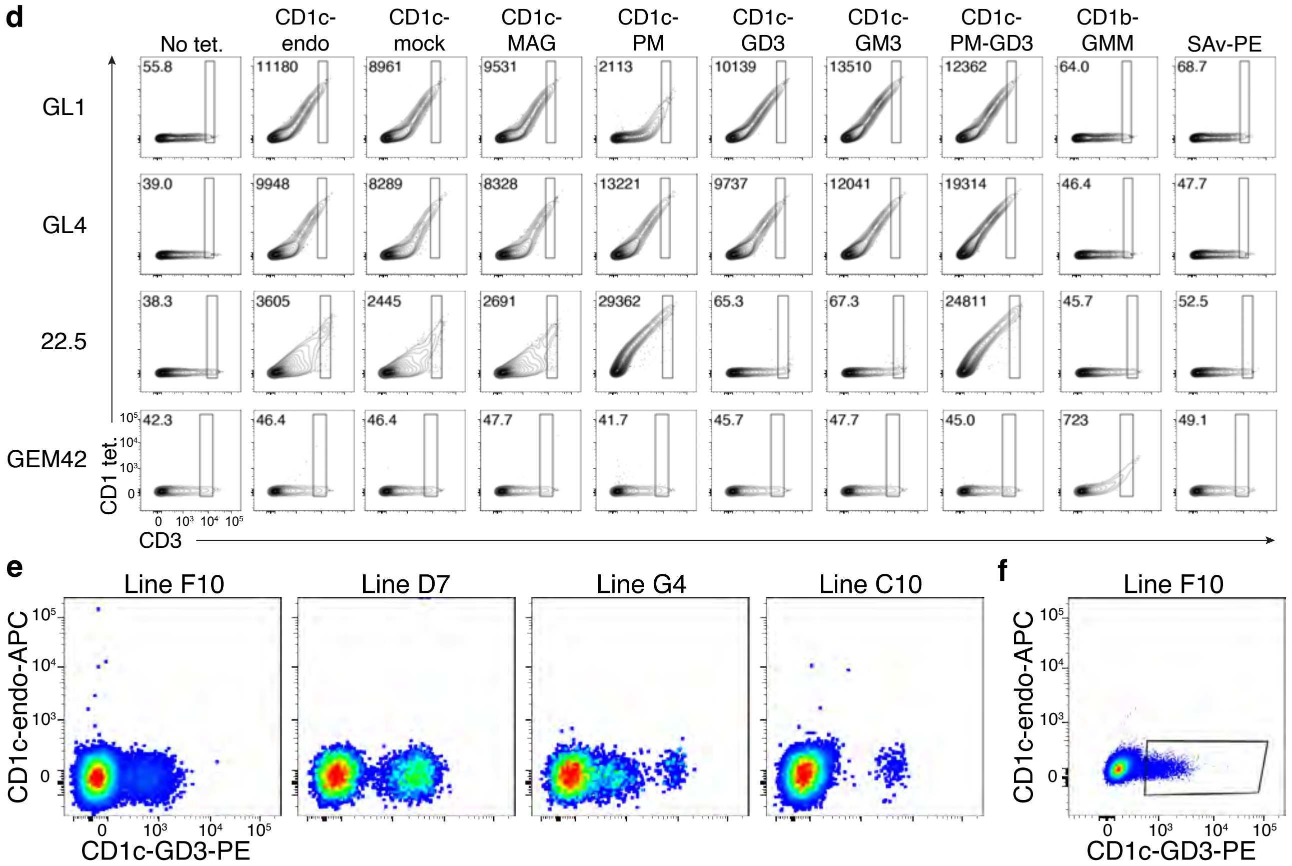


**Figure S8. Sequencing and identification of ganglioside reactive T cells**

**a.** Pie charts showing TCR gene usage of CD1c-restricted T cells that have higher avidity toward CD1c-GD3 than CD1c-endo. **b.** Sorting strategy of identification for GL1 and GL4 TCRs. Expanded T cells were stained with CD1c-GD3 tetramers and single-cell sorted for TCR sequencing. **c.** TCR sequences of clones GL1 and GL4 sorted from the polyclonal populations in panel **b**. **d.** The second independent repeat of **Figure 5c**. Contour plots showing CD1-lipid tetramer staining on HEK293T.SCARB1$^{-/-}$ cells transiently transfected to express CD1c reactive TCRs in the panel **b** or control TCRs, 22.5 (CD1c-PM reactive) or GEM42 (CD1b-GMM reactive). **e.** Expanded tonsillar T cells were dual stained with CD1c-GD3-PE and CD1c-endo-APC tetramers that exhibited minimal autoreactivity. **f.** Sorting strategy of establishing tonsillar T cell clone F10 from expanded T cells dual stained with CD1c-GD3-PE and CD1c-endo-APC tetramers. CD1c-GD3-PE tetramer$^{+}$ cells were sorted for another round of expansion.

### Table S1: Crystal data collection and refinement statistics

| | CD1c-GD3 | CD1c- GM3 | CD1c-MPM-GD3 | CD1c-MPM-GM1 | CD1c-mock | CD1c-PM | CD1a-GD3 | CD1b-GD3 |
|---|---|---|---|---|---|---|---|---|
| PDB ID | 9OHT | 9OHU | 9OHV | 9OHW | 9OHX | 9OHY | 9OHZ | 9OI0 |
| **Data collection** | | | | | | | | |
| Resolution (Å) | 43.64 – 2.98 (3.17 – 2.98) | 44.40 – 2.90 (3.08 – 2.90) | 47.32 – 2.12 (2.18 – 2.12) | 48.83 – 2.49 (2.59 – 2.49) | 47.87 – 1.68 (1.71 – 1.68) | 47.31 – 2.20 (2.27 – 2.20) | 46.18 – 2.14 (2.20 – 2.14) | 49.02 – 1.84 (1.94 – 1.84) |
| Space group | $P3_121$ | $P3_121$ | $P2_12_12_1$ | $P2_12_12_1$ | $P2_12_12_1$ | $P2_12_12_1$ | $P2_12_12_1$ | $P2_12_12_1$ |
| Cell dimensions a, b, c (Å) | 89.50, 89.50, 158.44 | 88.80, 88.80, 157.20 | 54.92, 84.94, 93.22 | 57.62, 86.56, 91.98 | 55.94, 84.66, 92.51 | 54.04, 83.47, 92.54 | 42.39, 90.35, 107.44 | 57.57, 77.85, 92.12 |
| Total reflections | 132394 (21947) | 165992 (27113) | 169752 (12862) | 104422 (11552) | 687806 (35258) | 247312 (21323) | 151229 (12411) | 1195235 (110294) |
| Unique reflections | 15589 (2467) | 16511 (2621) | 25243 (1929) | 16415 (1832) | 50885 (2571) | 22138 (1906) | 23537 (1885) | 36782 (5313) |
| Multiplicity | 8.5 (8.9) | 10.1 (10.3) | 6.7 (6.7) | 6.3 (6.2) | 13.5 (13.7) | 11.2 (11.2) | 6.4 (6.6) | 32.5 (20.8) |
| Completeness (%) | 100.0 (100.0) | 100.0 (100.0) | 99.5 (94.5) | 99.6 (100.0) | 100.0 (100.0) | 100.0 (100.0) | 100.0 (100.0) | 100.0 (100.0) |
| Mean I/s(I) | 10.8 (2.2) | 13.8 (2.5) | 10.6 (1.8) | 9.4 (2.5) | 18.2 (2.7) | 8.9 (2.3) | 11.8 (2.2) | 10.3 (2.0) |
| Rpim (%) | 6.0 (61.3) | 4.2 (56.7) | 3.9 (52.8) | 6.8 (68.9) | 2.2 (55.9) | 5.3 (43.6) | 3.8 (34.9) | 7.2 (41.5) |
| CC1/2 (%) | 99.6 (53.3) | 99.3 (65.3) | 99.7 (70.7) | 98.3 (54.5) | 99.9 (89.2) | 99.1 (85.3) | 99.8 (74.9) | 99.6 (75.7) |
| **Refinement** | | | | | | | | |
| Rwork (%) | 20.06 | 20.09 | 22.76 | 20.65 | 19.07 | 20.11 | 21.71 | 21.12 |
| Rfree (%) | 23.51 | 24.47 | 26.53 | 24.80 | 21.61 | 24.17 | 24.72 | 26.73 |
| R.m.s.d bond length (Å) | 0.011 | 0.004 | 0.009 | 0.008 | 0.006 | 0.008 | 0.006 | 0.010 |
| R.m.s.d bond angle (°) | 1.30 | 0.76 | 1.20 | 1.10 | 0.91 | 1.04 | 1.23 | 1.41 |
| Ramachandran plot | | | | | | | | |
| Favoured (%) | 95.73 | 95.48 | 95.95 | 95.20 | 98.92 | 95.66 | 94.55 | 98.67 |
| Allowed (%) | 4.27 | 3.72 | 3.78 | 3.73 | 1.08 | 3.79 | 3.81 | 1.33 |
| Outliers (%) | 0.00 | 0.80 | 0.27 | 1.07 | 0.00 | 0.54 | 1.63 | 0.00 |

*Values in parentheses are for highest-resolution shell

**Table S2: Primary sequence identity of CD1c between human and other species**

| Species | Scientific Name | Identity to human CD1c (%) |
|---|---|---|
| Alligator | *Alligator mississippiensis* | 28.35 |
| Panda | *Ailuropoda melanoleuca* | 47.91 |
| Gorilla | *Gorilla gorilla_gorilla* | 98.20 |
| Mangabey | *Cercocebus atys* | 90.49 |
| Monkey | *Chlorocebus sabaeus* | 90.69 |
| Rhesus macaque | *Macaca mulata* | 90.69 |
| Drill | *Mandrillus leucophaeus* | 90.69 |
| Angola colobus | *Colobus angolensis palliatus* | 91.25 |
| Elephant | *Loxodonta africana* | 69.07 |
| Fur seal | *Callorhinus ursinus* | 69.59 |
| Horseshoe bat | *Rhinolophus ferrumequinum* | 70.32 |
| Horse | *Equus caballus* | 69.35 |
| Guinea pig | *Cavia porcellus* | 55.26 |
| Goat | *Capra hircus* | 54.55 |
